# A unique C-terminal domain contributes to the molecular function of restorer-of-fertility proteins in plant mitochondria

**DOI:** 10.1101/2023.05.02.538898

**Authors:** Sang Dang Huynh, Joanna Melonek, Catherine Colas des Francs-Small, Charles S. Bond, Ian Small

## Abstract

*Restorer-of-fertility* (*Rf*) genes have practical applications in hybrid seed production as a means to control self-pollination. They encode pentatricopeptide repeat (PPR) proteins that are targeted to mitochondria where they specifically bind to transcripts that induce cytoplasmic male sterility and repress their expression. In searching for a molecular signature unique to this class of proteins, we found that a majority of known Rf proteins have a unique domain, which we called RfCTD (*Restorer-of-fertility C-terminal domain*), and its presence correlates with the ability to induce cleavage of the mitochondrial RNA target. We constructed a sequence profile that can quickly and accurately identify RfCTD sequences in plant genomes or transcriptomes. We screened 219 angiosperm genomes from 123 genera and found that each diploid genome encodes, on average, 25 Rf-like (RFL) proteins, of which approximately 55% contain the C-terminal signature domain. This screen also revealed considerable variation in RFL gene numbers across flowering plants. We observed that plant genera with bisexual flowers have significantly higher numbers of RFL genes compared to those with unisexual flowers, consistent with a role of these proteins in restoration of male fertility.

Finally, we show that removing the RfCTD from the RFL protein RNA PROCESSING FACTOR 2-*nad6* prevented cleavage of its RNA target, the *nad6* transcript, in *Arabidopsis thaliana* mitochondria. This research provides a simple way of identifying putative *Rf* candidates in genome sequences, new insights into the molecular mode of action of Rf proteins in plant mitochondria and expands our understanding of the evolution of fertility restoration in flowering plants.

## Introduction

### Cytoplasmic male sterility and restorer-of-fertility genes

Cytoplasmic male sterility (CMS), a mitochondrially-encoded trait that leads to a failure of the plant to produce fertile pollen, can spontaneously arise in natural plant populations or be generated artificially through interspecific crosses or protoplast fusions (Kaul 1988; Chase 2007). As CMS requires the affected plant to be pollinated by pollen from another individual, it can be used to facilitate outcrossing and hybridisation. Hybrid breeding methods based on CMS are popular as they are cheap, effective and relatively easy to control, once the requisite genotypes have been established. CMS-based hybrid seed technology requires a three-line system that includes a CMS line, a maintainer line and a restorer line (Chen and Liu 2014). The CMS line has male-sterility-inducing cytoplasm with a CMS-causing gene and lacks functional Restorer-of-fertility (*Rf)* genes (Chen and Liu 2014). Hence, this line is used as the female parent because it cannot produce functional pollen. In contrast to the CMS line, the restorer line has a functional *Rf* gene (or genes) located in the nuclear genome. *Rf* genes suppress the action of the corresponding CMS-causing genes, so the restorer line can act as the male parent in the crosses with the CMS line to create F_1_ hybrid seeds (Chen and Liu 2014). In the F_1_ plants, the *Rf* gene restores male fertility by blocking the expression of the CMS-causing gene in mitochondria. Identification of corresponding *Rf* and CMS genes is essential in hybrid seed production which requires full fertility restoration to ensure high seed-set.

### Rf proteins belong to the family of pentatricopeptide repeat (PPR) proteins

Most *Rf* genes belong to a specific clade (*Rf*-like, or RFL genes) of a much larger family of genes encoding RNA-binding organelle-targeted pentatricopeptide repeat (PPR) proteins that influence the processing, editing, splicing, stability and translation of specific organellar RNAs (Barkan and Small 2014). The PPR family is highly expanded in plants, comprising over 400 genes in most angiosperms (Lurin et al. 2004; Cheng et al. 2016). PPR proteins can be divided into two classes (P and PLS) based on the type of PPR motifs present (Schmitz-Linneweber and Small 2008). All functionally characterised PPR Rf proteins belong to the RFL clade of the P class (Fujii et al. 2011; Dahan and Mireau 2013; Melonek et al. 2016). One of the characteristic features of RFL genes is that they are located in clusters in a few genomic locations (Melonek et al. 2016, 2019; Melonek and Small 2022). These high-density clusters are the site of proliferation of new RFL gene variants, probably via unequal crossover events (Melonek et al. 2016). Like most other P class PPR proteins, RFL proteins appear to lack any catalytic domains and are thought to passively bind RNA (Kindgren et al. 2015; McDermott et al. 2018).

### Our knowledge of the mechanism underlying fertility restoration conferred by Rf proteins is limited

The majority of Rf proteins have been shown to be associated with RNA processing in plant mitochondria (Wang et al. 2006; Binder et al. 2013; Melonek et al. 2021; Jiang et al. 2022). Petunia Rf-PPR592 was shown to act in a post-transcriptional manner to reduce the amount of the CMS-associated PCF protein (Gillman et al. 2007), but how it did this remained unknown. The first insights on the possible mode of action of Rf proteins came from studies on CMS and fertility restoration in rice (Kazama and Toriyama 2003; Komori et al. 2004; Akagi et al. 2004). Here, two genes named *Rf1a* and *Rf1b* were found to restore rice BT-type CMS (Wang et al. 2006). Based on the molecular characterisation of the Rf1a and Rf1b proteins, the authors concluded that the Rf1a protein is able to induce cleavage of the *B-atp6/orf79* CMS-transcript at a specific site, whereas Rf1b causes degradation of the same transcript via an unknown mechanism (Wang et al. 2006). Similarly, the *OsRf19* gene, located in the same cluster of RFL genes on chromosome 10 in rice as *Rf1*, was shown to induce cleavage of the CMS-causing *fa182* transcript (Jiang et al. 2022). In bread wheat, two restorer proteins, Rf1 and Rf3, bind to their CMS-target, *orf279*, at two different sites and induce RNA cleavage in two different positions (Melonek et al. 2021).

Several of the 26 RFL proteins designated as RNA PROCESSING FACTORs (RPFs) in *Arabidopsis thaliana* were characterised to be required for efficient 5’ end maturation of mitochondrial mRNA and rRNA (Binder et al. 2013, 2016). For example, RPF2 is needed for maturation of the 5’ end of *nad9* and *cox3* mRNA (Jonietz et al. 2010) and RPF1 for that of *nad4* gene (Hölzle et al. 2011). Recent studies have shown that it is possible to re-target RPF2 protein from its natural targets to novel targets by modifying amino acids involved in binding to RNA (following the so-called ‘PPR-code’ (Barkan et al. 2012; Yagi et al. 2013; Yan et al. 2019) and induce RNA cleavage (Colas des Francs-Small et al. 2018; Yang et al. 2022). For example, modified versions of RPF2 were shown to bind and induce cleavage of *nad6* and *atp1* RNAs (Colas des Francs-Small et al. 2018; Yang et al. 2022). MNU1 and MNU2 endonucleases have been proposed as possible interaction partners of RPF2 to carry out the RNA cleavage (Stoll and Binder 2016), however these observations were not confirmed in studies with RPF2 targeted to a different site (Colas des Francs-Small et al. 2018). In addition, it has been reported that *Arabidopsis* RFL2 requires mitochondrial proteinaceous RNase P (PRORP) for endonucleolytic cleavage of the mitochondrial *orf291* transcript (Fujii et al. 2016). Although this study demonstrated that RFL2 protein binds directly to the *orf291* transcript, it is not clear how the RFL2 protein recruits PRORP. More recently, translational blockage rather than RNA cleavage has been proposed as a mode of action of Rfo (PPR-B) protein in *Brassica* plants carrying Ogura CMS (Wang et al. 2021a). Thus, despite these extensive studies on Rf and RFL proteins in *Arabidopsis* and other plant species, there is still no common view about the mechanism underlying the molecular function(s) of RFL proteins (Fujii et al. 2016; Binder et al. 2016; Melonek et al. 2021). A common (but not universal) observation is that RFL proteins cause cleavage or degradation of their target RNAs in mitochondria. As RNA binding by PPR proteins is more usually associated with stabilisation of RNAs (Pfalz et al. 2009; Prikryl et al. 2011; Ruwe and Schmitz-Linneweber 2012; Hammani et al. 2012), this implies that Rf proteins contain a specific feature that either directly or indirectly induces RNA cleavage, perhaps in association with other factors.

We started this work by looking for distinctive features unique to RFL proteins, and discovered one within the C-terminal region of many, but not all, RFL proteins. This C-terminal domain can be identified in candidate RFL proteins from virtually all flowering plants that we have looked at (164 species). We use this RFL marker to explore the broad phylogenetic distribution of RFL genes across flowering plants and discover strong correlations with reproduction strategy, probably linked to the role of the genes in suppressing naturally-occurring CMS. Some intriguing exceptions hint that these proteins may have been co-opted into different roles in some plant groups. To find out if this signature sequence has a functional role, we deleted this domain from RNA PROCESSING FACTOR 2-*nad6*, a cleavage-inducing *Arabidopsis* RFL, and found that this prevents cleavage (and suppression of expression) of its target transcript.

## MATERIALS AND METHODS

### Data used in this study

The genome sequences of 243 plant species were downloaded from *Phytozome13* (https://phytozome-next.jgi.doe.gov/) and other publicly available sequence databases as specified in Supplementary Table 2. The genomic sequences were screened for open reading frames (ORFs) in six-frame translations with the *getorf* program of the EMBOSS 6.6.0 package (Rice et al. 2000). Predicted ORFs were screened for the presence of P- and PLS-class PPR motifs using *hmmsearch* from the HMMER 3.2.1 package (Eddy 2011) (http://hmmer.org) and hidden Markov models defined by *hmmbuild* (Cheng et al. 2016). RFL proteins were identified via the OrthoMCL database (Fischer et al. 2011) (http://www.orthomcl.org/orthomcl/) hosted by the VEuPathDB Galaxy Data Analysis Service (https://galaxyproject.org/use/veupathdb/). The OrthoGroup containing RFL proteins was identified by the presence of reference RFL sequences and the highest number of proteins assigned to it among all OrthoGroups generated by OrthoMCL (Melonek et al. 2016). Alternatively, the P class PPR protein sequences were aligned with Muscle v3.8.425 (Edgar 2004) and a tree was generated using FastTree v2.1.11 (Price et al. 2010). RFL PPR sequences form a distinct clade in such trees as described previously (Melonek et al. 2016).

### Bioinformatic pipeline to build the RfCTD profile and screen for PPR sequences containing RfCTDs

464 RfCTD sequences identified in a preliminary screen of our PPR database (https://ppr.plantenergy.uwa.edu.au/) (Cheng et al. 2016) with the last 60 amino acids of *Arabidopsis thaliana* RPF2 and the last 71 amino acids of *Oryza sativa* were used to build an initial RfCTD profile with the *hmmbuild* program from the HMMER 3.2.1 package (Eddy 2011). This profile was then used to screen the P class PPR proteins found in the 243 plant genomes using the *hmmsearch program*. The identified RfCTDs were extracted, and redundant sequences were removed with CD-Hit v4.6.4 (Huang et al. 2010) with a threshold of 90%. *Hmmbuild* was used to create the final RfCTD profile from an alignment of 1,486 non-redundant sequences. The RfCTD profile was incorporated into PPRfinder code (Cheng et al. 2016) and the screen of the 243 genomes was repeated. Final filtering parameters of *hmmsearch* score >= 50, no overlapping with a PPR motif with a higher score and the RfCTD being the last motif in the protein were used to qualify a protein as containing RfCTD.

### Building an RfCTD sequence logo and comparing it to that of the last two PPR motifs

After reducing the redundancy by CD-hit at threshold of 90%, a sequence logo was built using the SKYLIGN web browser (http://skylign.org/) following the *information content - above background* setting to calculate letter- and stack-heights for each column of the alignment. To compare the RfCTD with P motifs, a corresponding logo of the two last P class motifs was also made from the same 1,486 PPR proteins. Sequence alignment was conducted with MAFFT v7.450 (Katoh and Standley 2013) in Geneious v2023.0.1 (https://www.geneious.com/) before extracting the two last PPR motifs. The logo of the last two PPR motifs were made using the same SKYLIGN web browser with the same settings.

### Biostatistical analysis for relationship between reproductive strategies and RfCTDs and RFLs

Reproductive morphologies (hermaphroditic, monoecious, dioecious, gynodioecious, and CMS) for each genus were obtained based on data from (Kaul 1988; Dufaÿ et al. 2014; Renner 2014)). Boxplots and dotplots of pRfCTD and pRFL across reproductive types were created with the Plots.jl package (https://docs.juliaplots.org/stable/) in Julia v1.8.4. As the pRfCTD and pRFL are not normally distributed, one-tailed Mann-Whitney U tests using dioecious taxa as reference were carried out with the HypothesisTests.jl package (https://juliastats.org/HypothesisTests.jl/stable/) in Julia v1.8.4 to test for significant differences in long RfCTD and long RFL frequency between different groups of taxa with different reproductive morphologies.

### Modelling the structure of the RfCTD

A structural model of *Arabidopsis thaliana* RPF2 protein was predicted by AlphaFold (Jumper et al. 2021) (https://alphafold.ebi.ac.uk/entry/Q9SXD1) with UniProt ID Q9SXD1. Per-residue confidence scores (pLDDT) for residues of the RfCTD were collected before calculating the average pLDDT for the domain. The created model was saved in PDB format and imported into PyMOL. In addition, a structural model of a synthetic P class PPR protein (4OZS) (Gully et al. 2015) was downloaded from Protein Data Bank (PDB) (https://www.rcsb.org/structure/4OZS).

### Plant material

Seeds of *Arabidopsis thaliana* ecotype Columbia (Col-0), *rpf2* mutant (Jonietz et al. 2010), *rpf2::RPF2nad6* complemented line *(Colas des Francs-Small et al. 2018)* and transformants generated within this study were grown on soil mixed with vermiculite and perlite in 3:1:1 ratio in long-day conditions with 16 hours light and 8 hours dark at 22°C in growth chambers.

### Plasmid cloning and Agrobacterium-mediated transformation of Col-0 plants

The truncated version of the RPF2*nad6* was generated by PCR amplification using Takara PrimeStar polymerase (TaKaRa) and a primer pair designed to remove the last 180 nucleotides of the RPF2*nad6* coding sequence representing the RfCTD sequence (Supplementary table 6). The amplified DNA fragment was gel-purified using QIAquick Gel Extraction Kit (Qiagen) and inserted into pDONR207 using Invitrogen Gateway® cloning technology following manufacturer’s guidelines (ThermoFisher Scientific). The obtained constructs were transformed into *rpf2* plants by floral dip with *Agrobacterium tumefaciens* (Shaw 1995). Seeds harvested approximately one month after the transformation were screened on Murashige & Skoog (MS) medium plates (PhytoTechnology Laboratories) supplemented with hygromycin B (PhytoTechnology Laboratories) at a final concentration of 50 µg/ml. The hygromycin B-resistant transformants were identified and the presence of the transgene was confirmed by genomic DNA extraction following the CTAB method (Doyle and Doyle 1987) and PCR with primer RPF2-2473 and 3FLAG-R2 (Supplementary Table 6). Expression of the transgene was verified by qRT-PCR analysis using Invitrogen SuperScript III Reverse transcriptase (ThermoFisher Scientific) sequence specific primer (Supplementary Table 6).

### Protein analysis and immunological detection

Arabidopsis leaves were ground into powder by mortar and pestle in liquid nitrogen. To extract proteins, the frozen plant material was resuspended in homogenisation buffer (0.0625 M Tris/HCl pH6.8, 1% (w/v) sodium dodecyl sulfate (SDS), 10% Glycerol, 0.01% 2-Mercaptoethanol) (Merck) and incubated at 95°C for 10 min following a previously published protocol (Dehesh et al. 1986). The protein samples were centrifuged at maximum speed for 1 min in an Eppendorf™ 5424 Microcentrifuge (Eppendorf). The proteins were separated on 10% SDS polyacrylamide gels and transferred onto Immuno-Blot PVDF membrane (BioRad) using Trans-Blot SD Semi-Dry Electrophoretic Transfer Cell (BioRad). Immunological detection was performed with anti-FLAG-tag antibody (ThermoFisher Scientific) diluted 1:1000 in 1xTBST buffer (50 mM Tris, 150 mM NaCl, 0.1% Tween 20) and the anti-mouse secondary antibody (Merck) diluted 1:10000 in 1xTBST. Chemiluminescent signal was detected with the Clarity™ Western ECL Blotting Substrate (BioRad) and images were taken using Amersham AI680 Western blot Imaging System (Cytvia).

### Analysis of RNA and Northern blot assay

Total RNA was extracted using the RNeasy Plant Mini Kit (Qiagen) and stored at -20°C. 10 µg of total RNA were separated on 1.2% agarose gels containing 10X MOPS buffer (3 M sodium acetate, 0.5 M EDTA, pH 7.0) and formaldehyde. RNA was transferred from the gel onto Amersham Hybond-N+ membrane (Cytvia) by capillary blotting with 10X Saline-Sodium Citrate (SSC) as transfer buffer for 16 hours at room temperature. Hybridisation with biotin-labelled short-DNA oligo as probe was conducted overnight at 50°C following published protocol (Colas des Francs-Small et al. 2018). Signal detection was performed using the Chemiluminescent Nucleic Acid Detection Module (ThermoFisher Scientific). Images were taken using Amersham AI680 Western blot Imaging System (Cytvia).

## RESULTS

### Identification of a C-terminal domain in Rf and RFL proteins that correlates with the ability to induce RN39A cleavage

Sequence alignment of all PPR Rf proteins identified to date (Supplementary Table 1) shows a 60–70 amino acid extension at their C-terminus after the last recognised PPR motif in most cases (Figure 1). The exceptions include the RsRfo, RsRfk and RsRf3 restorer proteins identified in radish (*Raphanus sativus*) and Rf1b in rice (Figure 1). Interestingly, these restorers also differ from most or all of the other known Rf or RPF proteins in that they do not appear to induce site-specific cleavage of their target RNA. Binding of RsRfo causes blockage of ribosome progression rather than induction of RNA cleavage (Wang et al. 2021a) and Rf1b, unlike Rf1a, appears to trigger RNA degradation without causing a specific cleavage event (Wang et al. 2006). Moreover, AtRFL8 that also lacks a C-terminal extension was shown not to induce RNA cleavage but rather act as a translational activator of the mitochondrial *ccmFN2* transcript (Nguyen et al. 2021). This apparent correlation between the C-terminal extensions and protein function triggered our interest to investigate these sequences in more detail.

**Figure 1.**
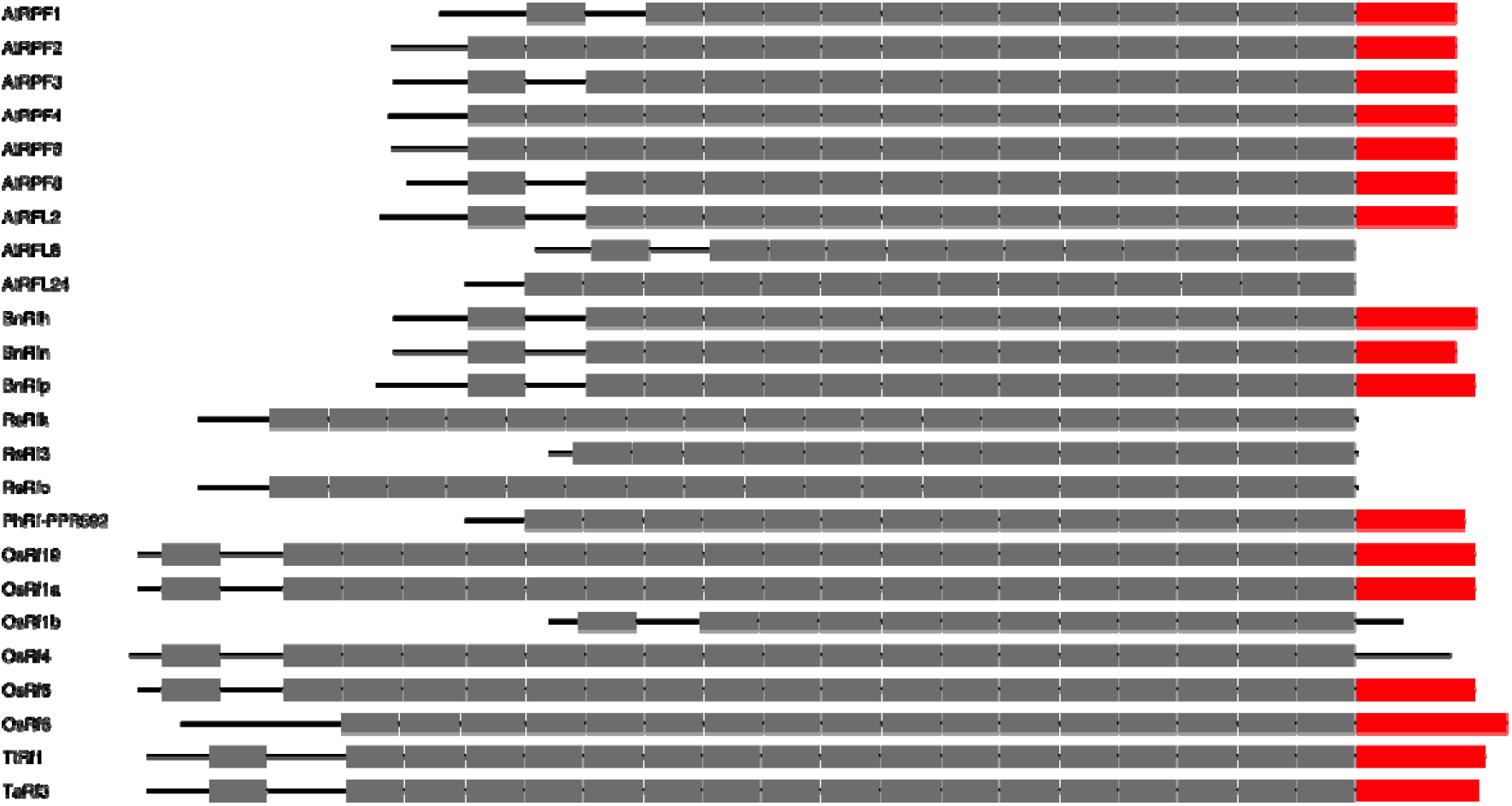
Presence of a 60–71 amino acid extension at the carboxy terminus of characterised Rf and RPF/RFL proteins. PPR motifs are shown as grey and the C-terminal extension as red boxes, respectively. The proteins are aligned at the end of the most C-terminal PPR motif, except for Rf1b which clearly corresponds to the N-terminal portion of Rf1a. AtRPF1–4, 6 and 8 are *Arabidopsis thaliana* RNA PROCESSING FACTORs (Jonietz et al. 2010, 2011; Hölzle et al. 2011; Stoll et al. 2015, 2017; Schleicher and Binder 2021); AtRFL2, 8 and 24 are *A. thaliana* RESTORER-OF-FERTILITY-LIKE proteins (Fujii et al. 2016; Durand et al. 2021; Nguyen et al. 2021); BnRfh restores fertility to *Brassica napus* plants carrying hau CMS (Wang et al. 2021b); BnRfn restores fertility to *B. napus* plants carrying nap CMS (Liu et al. 2017); BnRfp restores fertility to *B. napus* plants carrying pol CMS (Liu et al. 2016; Xiao et al. 2021); RsRfo restores fertility to *Raphanus sativus* and *Brassica* plants carrying ogu CMS (Brown et al. 2003); RsRfk restores fertility to *R. sativus* plants carrying kosena CMS (Koizuka et al. 2003); RsRf3 is *R. sativus* Restorer-of-fertility 3 (Wang et al. 2013); PhRf-PPR592 is *Petunia hybrida* Restorer-of-fertility PPR592 (Bentolila et al. 2002); OsRf1a–b (Wang et al. 2006), OsRf4–6 (Hu et al. 2012; Tang et al. 2014; Zhang et al. 2017) and OsRf19 (Jiang et al. 2022) are restorer-of-fertility proteins from *Oryza sativa*. TtRf1 and TaRf3 are restorer-of-fertility proteins from *Triticum timopheevii* and *T. aestivum* respectively (Melonek et al. 2021). Studies characterising the proteins shown in the figure alongside with their mitochondrial RNA targets, binding and cleavage sites (where known) are listed in Supplementary Table 1.

### Development of a sequence profile of the RfCTD

The last 60 amino acids of *A. thaliana* RPF2 and the last 71 amino acids of *O. sativa* Rf1a were used to search our plant PPR database (https://ppr.plantenergy.uwa.edu.au/) (Cheng et al. 2016). As a result, 464 similar C-terminal sequences were obtained, and they were employed to build an initial profile for searching for matching proteins in the proteomes of 243 plant genomes (Supplementary Table 2). A total of 5,665 matching sequences were identified and extracted. After reducing redundancy to a limit of 90% identity, a final CTD profile was constructed from the remaining 1,486 non-redundant sequences. In parallel, a sequence profile was constructed from the final two PPR motifs in the same 1,486 proteins. Figure 2 shows a comparison of the two sequence profiles.

**Figure 2.**
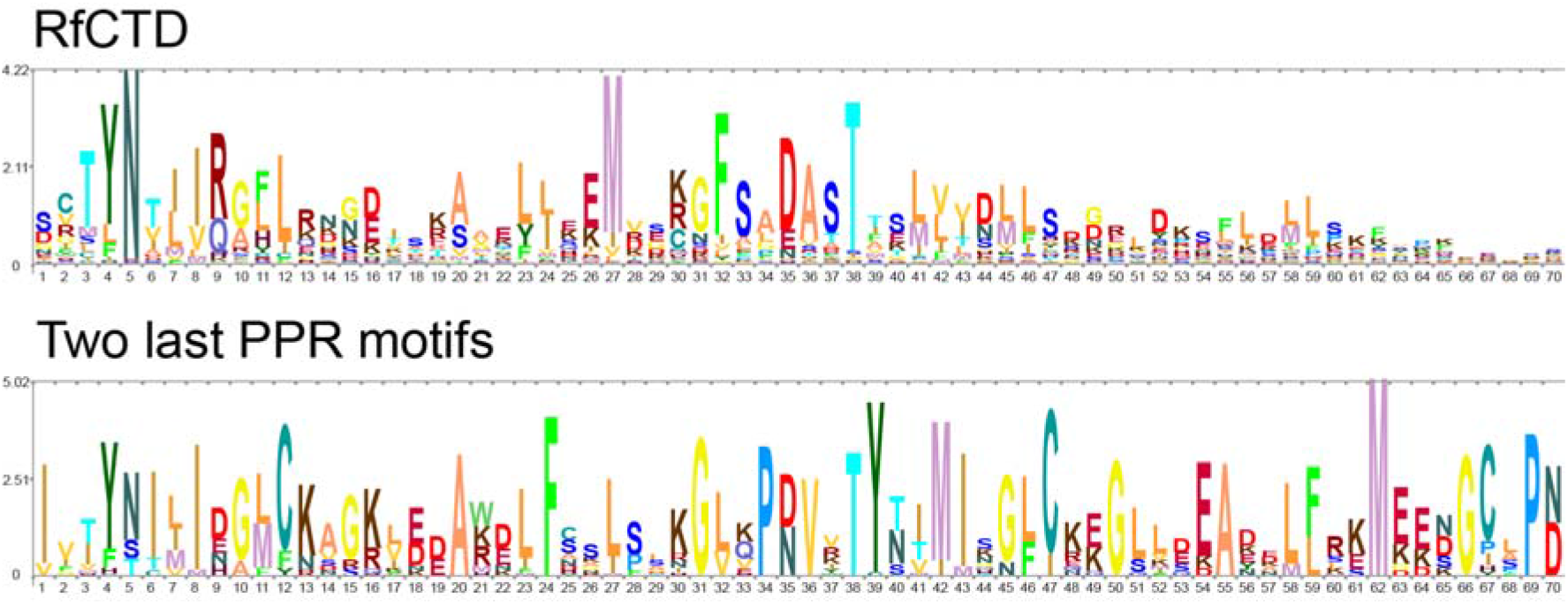
Sequence logos of the C-terminal extension (RfCTD) and two last PPR motifs generated using 1,486 non-redundant PPR sequences with SKYLIGN (http://skylign.org/).

The first 35 amino acids of the C-terminal sequences share some conserved residues with a typical P class PPR motif, e.g. amino acid residues 3-5 (threonine, tyrosine, asparagine), 31 (glycine), and 35 (aspartate); but there are other conserved amino acid residues that differ from a typical P class motif e.g. arginine at position 9, leucine at position 12, phenylalanine at position 32 and serine at position 33 that suggest that these sequences are not simply ‘degenerate’ PPR motifs. Beyond the conserved threonine at position 38 the C-terminal sequences are poorly conserved and differ considerably in length. The RfCTD profile detects all amino acid extensions present in the cloned Rf/RFL proteins that were identified during our initial analysis and are depicted in Figure 1.

We incorporated the RfCTD profile into PPRfinder (Cheng et al. 2016) and then used it to screen 243 proteomes obtained by translating entire genomes in all 6 frames. We focused on long PPR proteins (comprising at least 10 PPR motifs). As shown in Figure 3, high-scoring ‘hits’ are almost entirely at the C-terminus of PPR sequences, confirming the distinctness of these sequences. Based on these results we named this C-terminal extension Restorer-of-fertility C-terminal domain (RfCTD). Low-scoring matches to PPR motifs confirm that the RfCTD probably derived originally from PPR motifs and retains some similarity.

**Figure 3.**
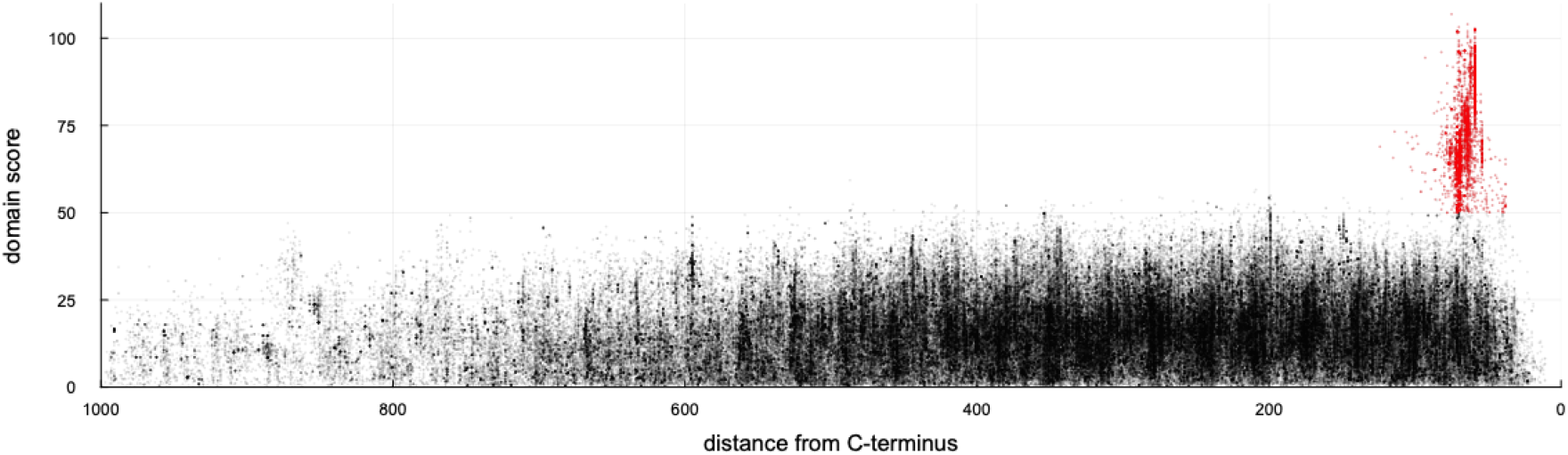
Screening long P class PPR proteins (>= 10 PPR motifs) for the presence of RfCTDs in 243 plant genomes. Each point represents one match between the RfCTD HMM and a protein sequence plotted according to the distance of the match from the C-terminus of the protein and the domain score calculated by hmmsearch. The matches depicted in red are those retained by PPRfinder (domain score >= 50 and higher than that of any overlapping PPR motif, C-terminal position); matches depicted in black are those discarded by PPRfinder. In total, 3,776 RfCTDs were identified.

### Variation of the number of RFLs in flowering plants

In total, 3,776 long PPR proteins containing RfCTDs were identified from 213 plant species representing 123 genera (Supplementary Table 3). There is a high degree of overlap between proteins identified as containing an RfCTD and RFLs identified by our usual method which relies on sequence clustering with OrthoMCL (Melonek et al. 2016) (Figure 4A). Ninety-seven percent (3,661) of the 3,776 RfCTDs identified in the screen are also identified as putative RFLs using OrthoMCL (Fischer et al. 2011) (Supplementary Table 3).

**Figure 4.**
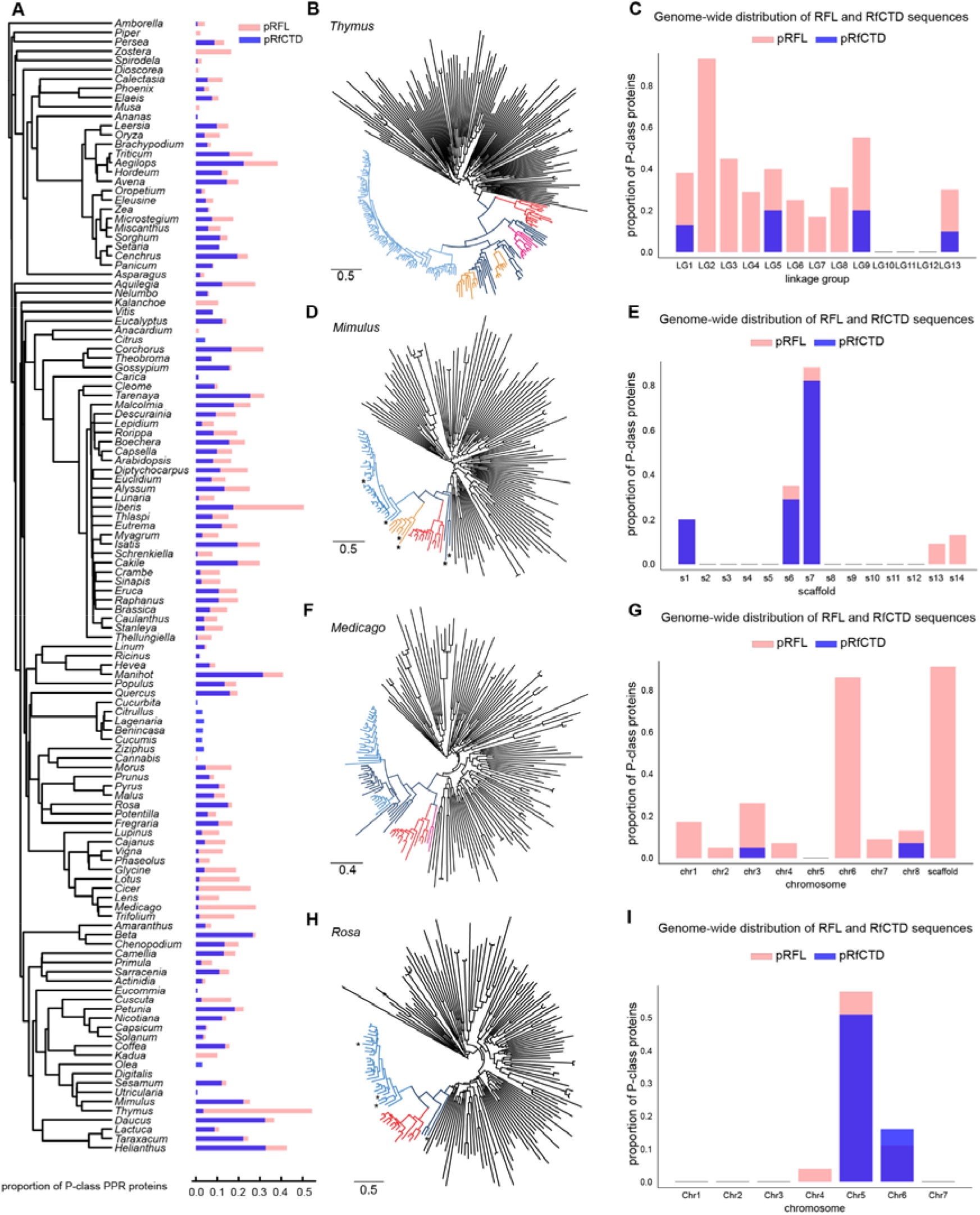
Identification of RFL and RfCTD sequences in 123 genera in flowering plants. **A**. Phylogenetic tree depicting the numbers of RFLs and RfCTDs in each genus. Blue bars indicate the number of RfCTDs detected using the *hmmer* profile developed in this study; pink bars indicate the number of RFLs predicted by OrthoMCL counts. In each case the counts are expressed as a proportion of the total number of P class PPR proteins to normalise for polyploidy and genome duplications. The tree was constructed with V.PhyloMaker2 (Jin and Qian 2022). **B**. Characterisation of the RFL and RfCTD sequences identified in thyme (*Thymus quinquecostatus).* P class PPR sequences were aligned with Muscle v3.8.425 (Edgar 2004) and the tree was generated using FastTree v2.1.11 (Price et al. 2010) in Geneious (https://www.geneious.com/). The midpoint rooting was calculated with iTOL and tree branches of the final tree were coloured also in iTOL v6.7.3 (https://itol.embl.de/) (Letunic and Bork 2021). The RFL clade (shown in blue) was identified through co-location with a reference set of 13 Arabidopsis RFL proteins (shown in red). RFL sequences located within the clusters on chromosomes 2 and 3 are coloured in light blue and orange, respectively. RFL sequences containing an RfCTD are shown in pink. **C**. Genome-wide distribution of RFL and RfCTDs in *Thymus*. The numbers are shown as the proportion of P class PPR proteins. **D**. Characterisation of the RFL and RfCTD sequences in *Mimulus guttatus.* RFL sequences located in a cluster on scaffolds 7 and 6 are coloured in light blue and orange, respectively. RFL sequences lacking RfCTD at their C-terminus are marked with an asterisk. **E**. Genome-wide distribution of RFL and RfCTDs in *Mimulus.* **F**. Characterisation of the RFL and RfCTD sequences in *Medicago truncatula.* RFL sequences located in a cluster on chromosome 6 are shown in light blue and RfCTDs are in pink. **G**. Genome-wide distribution of RFL and RfCTDs in *Medicago.* **H**. Characterisation of the RFL and RfCTD sequences in *Rosa chinensis*. The RFL clade located on chromosome 5 is shown in light blue. RFL sequences lacking RfCTD at their C-terminus are marked with an asterisk. **I**. Genome-wide distribution of RFL and RfCTDs in *Rosa*.

A small proportion (3%) consisting of 115 long proteins (>=10 PPR motifs) with RfCTD were not identified as RFL proteins by the OrthoMCL approach (Supplementary Table 3). Sixty-six of these are clustered in their respective genomes with RFL genes and thus are likely to be true RFL genes missed by OrthoMCL. In which case the adjusted number of long RfCTD proteins identified as RFLs would increase to 3,727 (98.7%). The remaining 49 are not clustered with other RFLs identified by OrthoMCL, or are in genomes lacking other RFLs, so these may be false positives. The comparison also suggests that only 54.9% of long RFL proteins (3,661 out of 6,659) contain an RfCTD. The proportion of known Rfs and RPFs with an RfCTD is 78.3%, a significantly higher proportion (Supplementary Table 1).

We conclude that the RfCTD profile is a rapid and easy way to identify putative Rf proteins, with a low false-positive rate if hits are filtered by PPRfinder. However, it cannot identify putative Rf proteins that lack the RfCTD, and nearly half of RFLs identified with the help of OrthoMCL lack RfCTDs (Supplementary Table 3).

The number of proteins containing an RfCTD varied from 0 to 125 across species (Supplementary Tables 2 and 4). Species belonging to *Cucurbitaceae, Eucommiaceae, Lentibulariaceae, Rubiaceae, Asparagaceae, Dioscoreaceae, Zosteraceae and Bromeliaceae* were found to contain distinctively small numbers of RfCTDs (less than 10) (Figure 4A, Supplementary Tables 2 and 4). OrthoMCL detected a low number of RFLs in these families too. In bread wheat (*Triticum aestivum),* a particularly high number of RfCTDs (from 109 to 125) was identified across different cultivars. Cassava (*Manihot esculenta*) also contained a high number of RfCTDs (∼100, depending on the cultivar). The number of RFLs predicted by RfCTD matches that predicted by OrthoMCL groups and generally covaries across angiosperms (with the OrthoMCL prediction being approximately two-fold higher on average). Outside angiosperms, though, the results differed greatly, with, for example, almost no RfCTDs detected in conifers but hundreds of RFLs predicted by OrthoMCL (Supplementary Table 3 and 4).

We have observed that, for some species, although OrthoMCL identified a high number of RFL genes, only a few of them appear to have RfCTDs. For example, in the recently published genome of five-ribbed thyme (*Thymus quinquecostatus)* (Sun et al. 2022), out of the 261 P class PPRs identified, 142 are RFLs according to OrthoMCL (Figure 4B and C; Supplementary Table 2). This amounts to a massive proportion, accounting for 54% of the total number of P class PPR proteins. The majority of RFL sequences are located in two clusters on chromosomes 2 and 3, containing 87 and 19 RFL genes, respectively (Figure 4C). Out of the 142 RFLs identified by OrthoMCL, only nine were found to have an RfCTD (Figure 4B and C). None of the RfCTDs were located within the RFL gene clusters on chromosomes 2 and 3, and most of them appeared as singlets (Figure 4C). In contrast, analysis of the RFL and RfCTD sequences in monkey flower (*Mimulus guttatus*), a close relative of thyme, revealed that the majority of RFLs in the species have an RfCTD (Figure 4D and E). Similarly, in legumes (family *Fabaceae*), a high number of RFL sequences identified by OrthoMCL, such as the 40–50 RFL sequences identified in the genome of *Medicago truncatula* (Figure 4F and G) *Lotus japonicus, Cicer arietinum* or *Glycine max* (Supplementary Figure 1), lack RfCTDs (Figure 4F and G). As in thyme, RFLs without the typical RfCTD are clustered in the genomes of legumes (Figure 4F, Supplementary Figure 2), while an average of two RfCTD sequences are present as singlets (Figure 4F; Supplementary Figure 1). On the other hand, in the genome of *Rosa chinensis*, from the sister group to legumes, the majority of identified RFLs have RfCTDs (Figure 4H and I).

The apparent absence of the conserved RfCTD in the majority of the RFLs identified in *Thymus* and legumes prompted us to investigate whether these proteins possess an additional PPR motif compared to proteins with an RfCTD (since the first half of the RfCTD is similar to a classical P class PPR motif). First, we calculated the average distance between the last PPR motif to the C-terminus, and found a shorter distance in *Thymus* compared to *Mimulus*, and in *Medicago* compared to *Rosa* (Supplementary Figure 3). Moreover, a global count of the number of PPR motifs per RFL protein showed that most plant RFL proteins have 13-14 PPR motifs, with the exception of grasses that have ∼18 PPR motifs (Supplementary Figure 3). However, except for *Thymus* and legumes where the non-RfCTD RFLs have one additional PPR motif on average, there is little or no variation in the number of P class PPR motifs between proteins containing an RfCTD and those without (Supplementary Figure 3C).

### Correlation between the number of RfCTDs and plant reproductive strategies

The survey of RfCTD sequences across angiosperms showed high variation between plant genera. As the RFL family is associated with fertility restoration in cytoplasmic male sterility systems in the few species where they have been studied, we tested whether these differences in RfCTD numbers might be linked to plant reproductive strategies. Figure 5A shows the number of RfCTDs in plant genera exhibiting hermaphroditism (bisexual flowers), monoecy (separate male and female flowers on the same plant), dioecy (male and female flowers on different plants) or gynodioecy (mixed populations of female and hermaphroditic plants). The figure also shows the numbers of RfCTDs in plant genera known to exhibit CMS (either spontaneously or following inter-specific crosses). For comparison, the number of RFL sequences is shown (Figure 5B).

**Figure 5.**
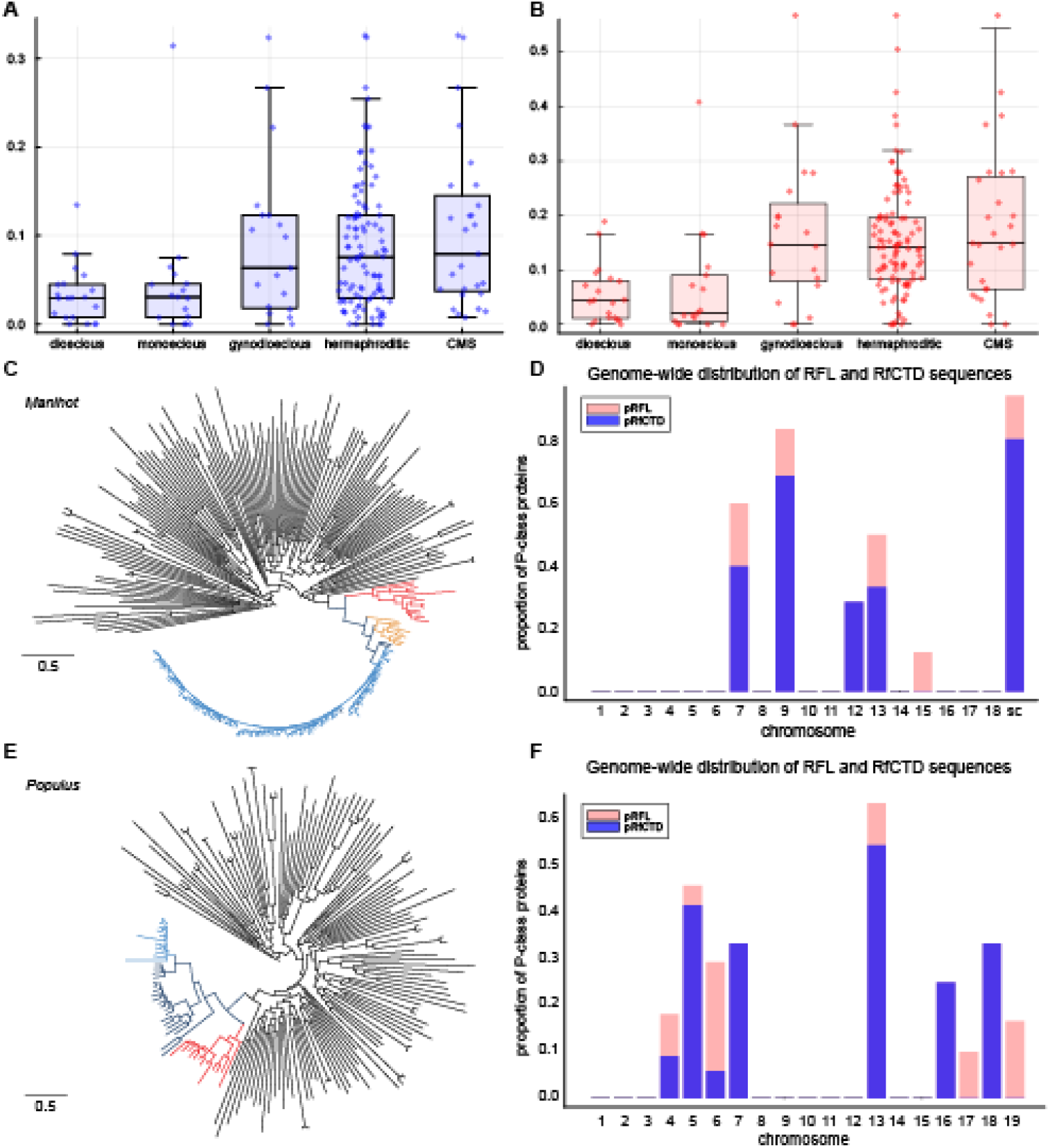
Correlation between proportion of long (>= 10 PPR motifs) RfCTD (pRfCTD) (**A**) and long (>=10 PPR motifs) RFL (pRFL) (**B**) sequences with flower morphology across 123 genera. **C**. Characterisation of the RFL clade in *Manihot esculenta.* P class PPR sequences were aligned with Muscle v3.8.425 (Edgar 2004) and the tree was generated using FastTree v2.1.11 (Price et al. 2010) in Geneious (https://www.geneious.com/). The midpoint rooting was calculated with iTOL and tree branches of the final tree were coloured also in iTOL v6.7.3 (https://itol.embl.de/) (Letunic and Bork 2021). The RFL clade was identified through co-location with a reference set of 13 Arabidopsis RFL proteins (shown in red). Sequences located within clusters on chromosomes 9 and 13 are shown in light blue and orange, respectively. **D**. Genome-wide distribution of RFL and RfCTDs in *Manihot*. The numbers are shown as the proportion of P class PPR proteins; sc indicates unanchored scaffolds. **E**. Characterisation of the RFL clade in *Populus trichocarpa*. Sequences clustered on chromosomes 5 and 13 are coloured in light blue and orange, respectively. **E**. Genome-wide distribution of RFL and RfCTDs in *Populus*.

*Morus notabilis* (dioecious) and cassava (*Manihot esculenta*, monoecious) were identified as outliers with proportion of long RfCTD (pRfCTD) to long RFL (pRFL) values of 0.15/0.19 and 0.36/0.41, respectively (Figure 5, Supplementary Table 4). In cassava, RFL genes are organised into a large cluster on chromosome 9 and a smaller cluster on chromosome 13 (Figure 5C). Many RFLs and RfCTDs were also found in poplar (*Populus trichocarpa*), another dioecious plant species (Figure 5; Supplementary Table 5). In this genome, the majority of RFL sequences are clustered on chromosomes 5 and 13 (Figure 5D).

In addition to analysing the genomes of core angiosperms, we also investigated the presence of RFLs and RfCTDs in the genomes of the representatives of the four major groups in gymnosperms (conifers, cycas, *Ginkgo, gnetophytes*) (Supplementary Table 2). We did not identify clades behaving like the RFL clade in the genomes of cycas and *Ginkgo biloba* (Supplementary Figure 4). In the genome of Chinese pine (*Pinus tabuliformis*) (Niu et al. 2022), 63 PPRs were predicted by OrthoMCL to be of RFL type. However, we did not find any RfCTD signature in these putative RFLs, and they did not group with the Arabidopsis RFLs in a phylogenetic tree (Supplementary Figure 5A). Forty-five of these putative RFLs clustered on chromosome 8. A combined tree with the P class PPR sequences from three other conifer genomes present in our dataset clearly shows that the sequences clustering on chromosome 8 are not closely related to RFL sequences found in angiosperms (Supplementary Figure 5B). These results suggest that some P class PPR sequences located in large genomic clusters can be found in conifer genomes, but these are unlikely to be closely related to angiosperm RFLs.

Within the flowering plants, we analysed the genomes of early-diverging angiosperms, the Amborellales (A) and Nymphaeales (N) orders (at the time of this study, there are no available reference genomes of Austrobaileyales (A) order). In the phylogenetic trees of P class PPR sequences from *Amborella trichopoda* and *Nymphaea colorata*, we did not find clades showing sequence similarity to the clade containing the reference set of Arabidopsis RFL sequences (Supplementary Figure 6). We do not find any clades that show evolutionary patterns typical of RFL sequences in the two genomes. Only one RfCTD and six putative RFLs were found in the genome of *A. trichopoda* (Supplementary Figure 6), however as the analysed genome is assembled only into short scaffolds, it is impossible to determine whether these cluster in the same genomic position. In contrast, in *Chloranthales,* an order among the early-diverging lineages of angiosperms, several P class PPR sequences were found to form a small clade in the phylogenetic tree (Supplementary Figure 6). Eight out of 9 sequences building the clade are located in a genomic cluster on chromosome 7 (Supplementary Table 3) and the majority of them were predicted as RFLs by OrthoMCL. Due to the close evolutionary relationships between Chloranthales and Magnoliids (Ma et al. 2021), we tested whether these sequences group with RFL sequences identified in avocado (*Persea americana* var. Hass), a representative of the Magnoliid clade, and they do (Supplementary Figure 6).

### Prediction of RfCTD structure

Next we used AlphaFold (Jumper et al. 2021) to obtain the structure of the RfCTD domain and compared it with the structure of a P class PPR motif. In a typical P class PPR motif, 35 amino acids form two α-helices (Figure 7). Base-specific RNA binding is based on the residues at position 5 in the first helix and the last residue in the loop between helices, i.e. position 35 in a classical P class PPR motif. The structure of the RfCTD of *A. thaliana* RPF2, predicted using AlphaFold (Jumper et al. 2021), consists of three long α-helices comprising 14 amino acids and one short α-helix consisting of 6 amino acids (Figure 7). The comparison between the structure of RfCTD structure and that of a synthetic P class PPR protein (4OZS) designed from consensus sequences (Gully et al. 2015) showed that the first two α helices of the RfCTD overlap with the two α helices of P motif while the third and fourth helices of the RfCTD do not overlap with two α helices of the next P motif (Figure 7B and C). The third helix was found to be displaced inward. The unusual arginine (R) at the 9^th^ position and leucine (L) at the 12^th^ position of the RfCTD may be linked to this structural divergence (Figure 7C). The predicted difference in the structure of the RfCTD compared to a typical P motif and the presence of unusual residues could infer that the RfCTD has a distinct function.

**Figure 7.**
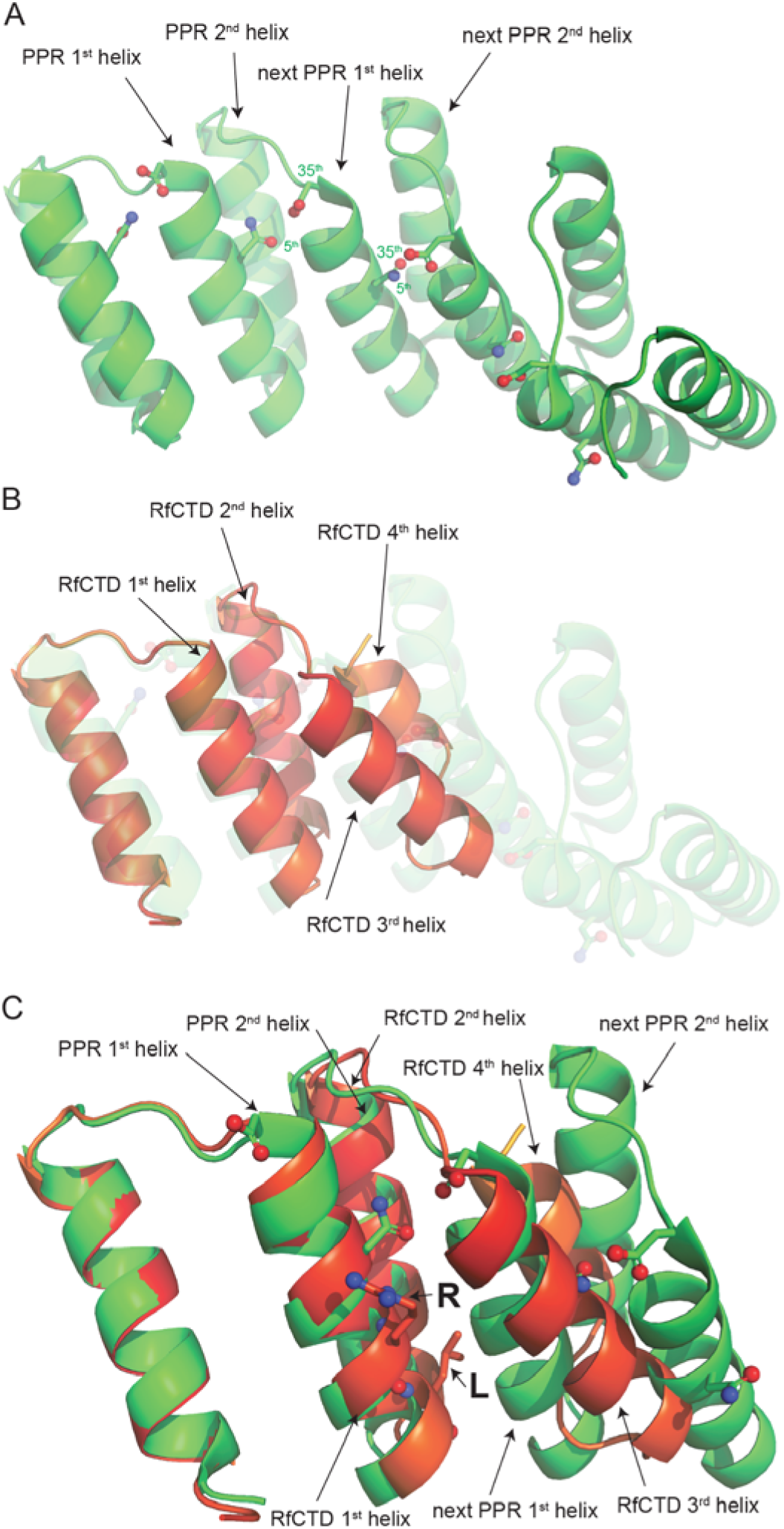
Comparison of the structure of RfCTD of *A. thaliana* RPF2 protein and that of a synthetic P class PPR protein. **A**. Structure model of 4OZS, a synthetic P class PPR protein (Gully *et al.*, 2015) (https://www.rcsb.org/structure/4OZS). Five PPR motifs, each folded into two α-helices are shown. The residues at positions 5 and 35 in the motif are shown. **B**. Structure model of RfCTD of *A. thaliana* RPF2 (red) overlaid on the shadow of the 4OZS P class PPR protein (light green). The first two α-helices of RfCTD overlap with two α-helices of a P class PPR motif while the 3^rd^ and 4^th^ helices of RfCTD do not overlap well with the equivalent PPR helices. **C**. The unusual residues of arginine (R) (position 9) and leucine (L) (position 12) in the RfCTD helix 2 are shown.

### Deletion of the RfCTD from RNA PROCESSING FACTOR 2-nad6 prevents cleavage of its target RNA

To test whether the RfCTD contributes to the biological function of RFL proteins, a truncated version of *RNA PROCESSING FACTOR 2-nad6* (RPF2*nad6*) was generated. The native sequence of RPF2 protein binds to the 5’ UTR of *cox3* and *nad9* mRNA (Jonietz et al. 2010), whereas the redesigned version RPF2*nad6* binds to the coding region of *nad6* mRNA and induces its cleavage (Colas des Francs-Small et al. 2018) (Figure 8A). To investigate the role of the RfCTD in the function of RPF2*nad6*, a truncated version of the protein was created by removing all 60 amino acids of the RfCTD (Figure 8A). To enable immunological detection of the truncated protein, a 3xFLAG tag was added directly after the last PPR motif (Figure 8A). The construct encoding RfCTD-truncated RPF2*nad6*-3xFLAG protein was transformed into Arabidopsis Col-0 background. In contrast to transgenic plants expressing full-length RPF2*nad6* that showed curled leaves, the transgenic plants expressing the RfCTD-truncated RPF2*nad6* did not show any visible phenotype and did not differ from Col- 0 or *rpf2* mutant plants (Figure 8B). Even though the accumulation of the truncated RPF2*nad6* protein in the transformants was confirmed immunologically (Figure 8C), the RPF2*nad6* protein lacking RfCTD was not biologically functional, as no cleavage in the *nad6* mRNA was observed in the Col-0 transformants (Figure 8D). Only the full-length *nad6* mRNA of ∼ 618 nucleotides was detected by northern blot (Figure 8D). Similarly, the same full-length *nad6* mRNA was detected in Col-0 and *rpf2* mutant plants lacking RPF2*nad6* protein (Figure 8D). In contrast, two transcripts corresponding to full-length and cleaved *nad6* mRNA were detected in the transformants expressing full-length RPF2*nad6* (containing RfCTD). This indicates that the RfCTD is essential for RPF2*nad6* to induce cleavage.

**Figure 8.**
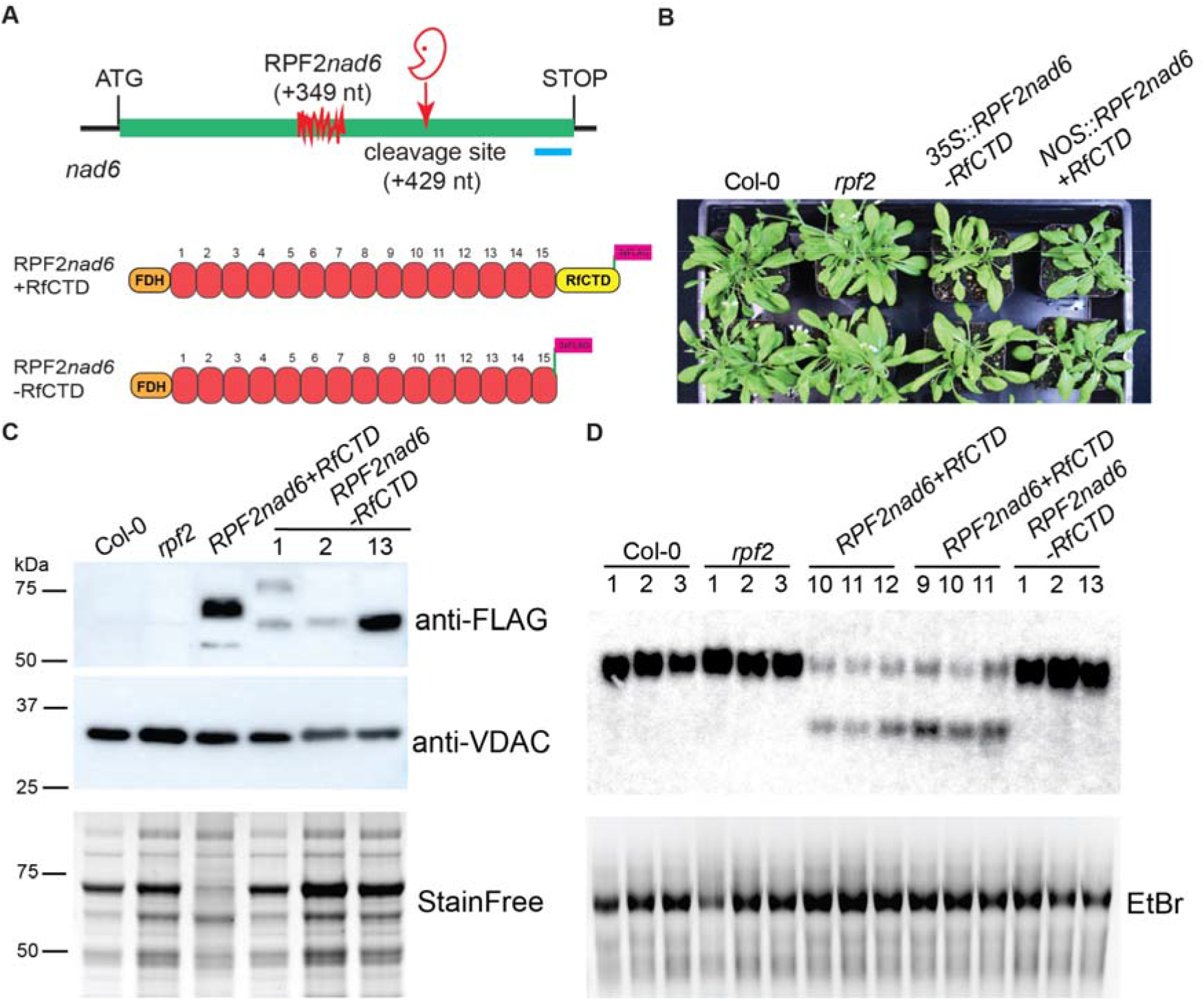
Contribution of the RfCTD to the biological function of RPF2*nad6* protein. **A**. Overview of the binding site of RPF2*nad6* and the induced cleavage site in the *nad6* transcript based on previously published data (Colas des Francs-Small et al. 2018). Schematic depiction of the full length *RPF2nad6* (RPF2*nad6*+RfCTD) and the truncated version of the protein (RPF2*nad6*-RfCTD) showing the PPR motifs as red boxes. FDH indicates the mitochondrial targeting sequence of formate dehydrogenase. Binding position of the 556 AS probe in the *nad6* gene, designed previously (Colas des Francs-Small et al. 2018), is shown as a blue bar. **B**. Phenotype of *Arabidopsis rpf2* plants transformed with construct encoding RfCTD-truncated (*rpf2::RPF2nad6-RfCTD*) and full-length (*rpf2::RPF2nad6*) RPF2*nad6* protein compared to *Arabidopsis* ecotype Col-0 and *rpf2* mutant. **C**. Immunological detection of the truncated and full-length RPF2*nad6* protein. Mitochondrial proteins from three independent lines were tested by western blot using anti-FLAG-tag antibody. The expected size of the RfCTD-truncated RPF2*nad6* protein fused with 3xFLAG is 64.7 kDa and that of full length RPF2*nad6* is 70 kDa. Mitochondrial proteins from Col-0 and *rpf2* mutants were used as negative controls. An antibody against VOLTAGE-DEPENDENT ANION CHANNEL 1 (VDAC1) protein was used as a reference for the presence of mitochondrial proteins in the samples. The size of the VDAC1 protein is estimated at 29 kDa. A stain-free SDS-PAGE gel is shown as reference for protein loading. M - protein molecular weight marker. **D**. Northern blot analysis of the *nad6* transcript in the RfCTD-truncated RPF2*nad6* transformants. Total RNA extracted from three independent plants of Arabidopsis ecotype Col-0, *rpf2*, and *rpf2* mutants transformed with *rpf2::RPF2nad6* or *rpf2::RPF2nad6-RfCTD* constructs, respectively, was analysed using probe 556 AS. A picture of the RNA gel stained with ethidium bromide (EtBr) is shown as a loading control.

## DISCUSSION

### Identification of a novel C-terminal domain in RFL proteins

In this study, we characterised a previously unrecognised domain, the RfCTD, characteristic of proteins in the RFL clade of P class PPR proteins. The RfCTD can be added to a list of other C-terminal domains linked to P class PPR proteins (Manna 2015), including a metallonuclease domain in members of PRORP proteins (Holzmann et al. 2008; Pinker et al. 2013), LAGLIDADG motifs in the splicing factor OTP51 (de Longevialle et al. 2008), an SMR domain in all members of the PPR-SMR group (Zoschke et al. 2013, 2016; Wu et al. 2016; Zhou et al. 2017; Takahashi et al. 2021), and the tRNA guanine-N7 methyltransferase (TGN) domain in the PPR-TGM subfamily (Manna and Barth 2013). While these C-terminal domains were most likely acquired by gene fusions (Liu et al. 2013; Manna and Barth 2013; Lechner et al. 2015; Gobert et al. 2019), in contrast, the origin of the RfCTD is different. The RfCTD and P motifs share conserved residues. This suggests that the RfCTD is derived from PPR motifs by sequence divergence, not by gene fusion.

### Correlation of numbers of RFLs/RfCTDs with reproductive strategies

Our analysis shows that hermaphrodite species with bisexual flowers tend to have a higher proportion of RfCTDs and RFLs compared to monoecious and dioecious species with unisexual flowers. The rapid evolution of RFL genes has been proposed to be driven by an ‘arms-race’ between mitochondrial and nuclear genomes (Touzet and Budar 2004). Once CMS occurs in a population to destroy male function, RFLs tend to be selected to control the expression of CMS to restore male function (Hanson and Bentolila 2004; Touzet and Budar 2004; Melonek et al. 2016). CMS naturally occurs in many hermaphroditic species where it is one of the possible causes of gynodioecy (Hanson 1991; Budar et al. 2003; Touzet and Budar 2004; Spigler and Ashman 2011; Yamauchi et al. 2019; Toriyama 2021). Greater numbers of RFLs in hermaphroditic and gynodioecious species found in our study tend to support the ‘arms-race’ proposal of coevolution of CMS and RFLs. Dioecious taxa contain significantly lower numbers of RFL (and RfCTD) proteins than CMS, gynodioecious and hermaphroditic taxa. The high number of RFLs in the latter taxa could be the result of RFL propagation driven by unequal crossover events (Melonek et al. 2016).

In vegetable and cereal crops, the many RFLs present in the genomes are often accompanied by several CMS sources documented in the species. For example, we found a large number of RFLs (77) in the carrot (*Daucus carota*) genome. The majority of the genes are clustered on chromosome 9 and encode proteins containing an RfCTD (69) (Supplementary Tables 3 and 4). So far, three types of CMS have been discovered in the species, and *Rf1*, the main restorer gene for petaloid CMS, the most widely used CMS in carrot hybrid breeding, has been mapped to chromosome 9 (Alessandro et al. 2013).

Most of the 98 RFL genes identified in the sunflower (*Helianthus annuus*) genome are located within three big clusters on chromosomes 3, 7 and 13, and have RfCTDs (Supplementary Table 3). The genomic locations of the RFL clusters coincide with the mapped positions of multiple restorer genes reported to be involved in fertility restoration of the PET1 and PET2 CMS types in the species (Sajer et al. 2020). This includes *Rf1*, *Rf-PET2* and *Rf5* mapped on chromosome 13, *Rf3* and *Msc1* on chromosome 7 and *Rf4* on chromosome 3, respectively (Sajer et al. 2020). Similar to carrot, the genomic regions carrying restorer genes in cereals, such as wheat, barley, and rye, overlap with the locations of RFL clusters (Melonek and Small 2022). These crops contain numerous RFLs and multiple CMS sources. In the Triticeae tribe, clusters of RFLs located on chromosome 1 and 6 (or 4 in rye) are frequent sources of new restorer gene variants (Melonek et al. 2019; Rabanus-Wallace et al. 2021). Approximately 50% of the RFL genes found in those genomes contain an RfCTD (Supplementary Table 3).

Also in *Thymus,* a gynodioecious species, a large number of RFLs coincides with the fact that CMS is very common in natural populations in the genus (Manicacci et al. 1996; Budar and Pelletier 2001). At least four types of CMS were described in garden thyme (*T. vulgaris*), which appear to be restored by distinct *Rf* loci (Budar et al. 2003). One could assume that a large reservoir of RFL genes in the genome helps generate new *Rf* gene variants to target novel *orfs* created in the mitochondrial genome. In favour of this hypothesis, an unusually high polymorphism of mitochondrial DNA has been reported in the species (Belhassen et al. 1993; Budar and Pelletier 2001). Interestingly, only nine of the 142 RFLs identified in the *T. quinquecostatus* genome contain an RfCTD (Figure 4; Supplementary Table 2 and 3). Further studies will be needed to determine whether *Thymus Rf* genes have developed a different mechanism of action than cleavage induction.

Despite a well-documented CMS in common bean (*Phaseolus vulgaris*) (Hervieu et al. 1993), only a relatively low number of RFLs (9) were found in the species (Supplementary Figure 1). This observation could be partially explained by the fact that the *Fr* gene reported to restore fertility of the CMS-Sprite in *P. vulgaris* works in a very different way to all other Rf genes by inducing physical loss of the CMS-inducing DNA region from the mitochondrial genome. The *pvs* region is located on an independently replicating subgenomic DNA molecule(Mackenzie and Chase 1990), therefore the *Fr* gene was proposed to trigger substoichiometric shifting in bean mitochondria, a phenomenon responsible for spontaneous reversion to fertility in cytoplasmic male sterile crop plants (Mackenzie and Chase 1990). As RFLs are RNA binding proteins, the *Fr* locus most likely does not encode a PPR protein.

The low values of RFL (and RfCTD) of dioecious taxa can be explained as CMS has no possible selective advantage in dioecious taxa (Wade and Goodnight 2006). These findings are consistent with reproductive mechanisms affecting the selection of CMS and *Rf* alleles (Fujii et al. 2011). Interestingly, CMS presents in some monoecious species, e.g., *Zea*, *Cucurbita*, and *Ricinus*, but both RFL and RfCTD values are generally significantly lower in monoecious taxa than in hermaphroditic taxa and are indeed similar to those of dioecious taxa, a surprising observation which requires further investigation.

### Possible other functions of RFL proteins than suppression of CMS

OrthoMCL and phylogenetic analyses revealed a high number (141) of RFL sequences in the *Manihot esculenta* (cassava-7) genome. RFLs constitute 55% of the total number of P class PPR sequences (255) identified by PPRfinder in the species. The two largest RFL gene clusters are located on chromosomes 9 and 13, comprising 99 and 14 RFL genes, respectively. 94 of these sequences contain the RfCTD sequence. In our analysis of the number of RFL/RfCTD and plant reproduction strategy, cassava is a clear outlier in monoecious species as its RfCTD and RFL proportion is much higher than the average. So far, no CMS has been reported in cassava, and the information about RFL and PPR proteins in this species is very limited. The reason for the high RFL numbers in cassava is unclear. Similarly, with an unexpectedly high proportion of RfCTD and RFLs, *Populus* is an outlier amongst dioecious taxa. The presence of many RFL genes in *Populus trichocarpa* (a strictly dioecious species) was noted previously (Fujii et al. 2011). Apart from *P. trichocarpa,* we also detected the presence of many RFLs in *P. alba* var*. pyramidalis* and *P. deltoides.* The high proportion of RFL proteins in the *Populus* genus strongly supports the idea that RFL gene propagation may sometimes be driven by other functional constraints than the necessity to suppress CMS (Dahan and Mireau 2013). For example, PPRs in poplar were reported to respond to biotic and abiotic stresses (Xing et al. 2018).

### Emergence of RFL sequences during plant evolution

In our studies of the few available chromosome-level assemblies of gymnosperm genomes, we did not identify PPR sequences similar to the RFL sequences found in core angiosperms. RFLs appear to be absent also in early-branching angiosperms, the Amborellales and Nymphaeales. While the absence of RFLs in Amborella, a dioecious species, is not surprising, their absence in Nymphaeales, a bisexual species, is more unusual and suggests that RFLs emerged only in euangiosperms following their divergence from the ANA grade (Anger et al. 2017). In the future, as more high-quality genome assemblies become available, their studies will provide a clearer picture of the RFL evolution in plants.

### The RfCTD is essential for the proper function of RFL proteins

Structural predictions of the AtRPF2 RfCTD reveal that its 3^rd^ and 4^th^ helices move inward and do not align with typical P motifs. In addition, the arginine (position 9) and leucine (position 12) conserved in RfCTDs are unusual residues in P motifs. These structural and sequence differences imply that this domain has a function distinct from the PPR motifs in AtRPF2.

The truncation experiments removing the RfCTD from RPF2*nad6* show that it is essential for the function of these proteins in *A. thaliana*. However, how the RfCTD contributes to the AtRPF2 function remains unclear. As the accumulation of truncated RPF2*nad6* was observed in mitochondrial extracts, it is unlikely that the RPF2 RfCTD is needed for expression of the RPF2 gene, the import of RPF2 into mitochondria, or the stability of the protein once inside. The RfCTD could play a role in helping the RFL proteins to bind to their targets, either by contributing directly to binding or by favouring correct folding of the rest of the protein. However, this seems unlikely as several known Rfs (e.g. Rfo, (Wang et al. 2021a) lack an RfCTD and yet still bind specifically to their target RNA. Rfo is currently the only example that demonstrates an Rf can cause ribosome blockage when it binds to the coding region of its target (Wang et al. 2021a). In the case of truncated RPF2*nad6*, effective ribosome blockage is unlikely because the transgenic plants did not have the same phenotype as transgenic plants containing RPF2*nad6* with RfCTD.

A second possible role of the RfCTD is to recruit an endoribonuclease to cleave the target RNA. This is our favoured hypothesis, as it fits with the observations that natural Rf proteins with the RfCTD domain are associated with RNA cleavage, whereas those that lack this domain are not. Knowledge about the interaction between RFLs and other partners is limited, but RFL2 is presumed to associate with the endonuclease PRORP1 (Stoll and Binder 2016; Fujii et al. 2016), and more recently PRORP1 was reported to directly interact with MNU2 via a specific domain of MNU2 and a proline-rich motif of PRORP1 (Bouchoucha et al. 2019).

### Potential future applications of the RfCTD profile

In breeding for better grain yield using CMS/*Rf* systems, it is vital to identify and use *Rf* genes capable of completely suppressing male sterility to recover as much fertility as possible in the F1 generation. We found that almost all PPR proteins that contain the RfCTD are RFL proteins. In addition, the presence of the RfCTD is correlated with the ability of Rf and RFL proteins to induce cleavage of their targets. Therefore, the RfCTD is a useful marker of RFLs that are more likely to be effective Rfs. Using *hmmsearch* with the RfCTD profile, we could check for the presence of the RfCTD in a large PPR data set from 243 plant genomes in a short time (a few minutes) compared to almost a week for identifying RFLs using OrthoMCL with the same database. Hence, this will help accelerate the development of hybrid breeding systems in crops where CMS/*Rf* systems are used. The number of sequenced crop genomes is rapidly increasing, creating an opportunity for identifying more functional *Rf* genes.

## ACKNOWLEDGEMENTS

SDH, JM, CCdF-S and IS received support from the Australian Research Council Centre (ARC) of Excellence in Plant Energy Biology (CE140100008) and the ARC Linkage grant LP200100547. SDH’s PhD studies were funded through the Australian Government International Research Training Program scholarship, The University of Western Australia Postgraduate Award, and Charles and Annie Neumann PhD in Agriculture top-up scholarship. This work was also supported by resources provided by the Pawsey Supercomputing Centre with funding from the Australian Government and the Government of Western Australia.

## AUTHOR CONTRIBUTIONS

JM and IS designed the study. SDH, JM, and IS performed the bioinformatic analyses. SDH and CCdF-S performed the molecular biology experiments. SDH and CB carried out protein structure predictions. JM and IS wrote the manuscript with input from all co-authors.

## DATA AVAILABILITY STATEMENT

The datasets generated during this study are available from DRYAD repository (https://doi.org/10.5061/dryad.2ngf1vhtq). PPRfinder can be downloaded from GitHub (https://github.com/ian-small).

## Supplementary Material

**Supplementary Figure 1.** Characterisation of the RFL clade in *Cicer arietinum*, *Glycine max*, *Lotus japonicus*, *Phaseolus vulgaris*.

**Supplementary Figure 2.** Distance of the last PPR motif from the end of the protein in *Thymus quinquecostatus, Mimulus guttatus, Medicago truncatula, Rosa chinensis*.

**Supplementary Figure 3.** Number of PPR motifs in RFL proteins across 243 genome data sets.

**Supplementary Figure 4.** Characterisation of the P class subfamily in gymnosperms.

**Supplementary Figure 5.** Evolutionary relationships among P class PPR proteins in conifers.

**Supplementary Figure 6.** Characterisation of the P class subfamily in basal angiosperms (*Amborella trichopoda, Nymphaea colorata)* (ANA grade) *and Chloranthus sessilifolius*.

**Supplementary Table 1.** Published Rf and RPF proteins.

**Supplementary Table 2.** Genome sequences used in the study.

**Supplementary Table 3.** Identified long (>=10 PPR motifs) P class, RFL and RfCTD sequences.

**Supplementary Table 4.** Proportions between long (>=10 PPR motifs) RFL and RfCTD sequences.

**Supplementary Table 5.** Plant reproductive strategies and presence/absence of CMS.

**Supplementary Table 6.** Oligo sequences used in this study.

