## Supplementary Figures for "A unique C-terminal domain contributes to the molecular function of restorer-of-fertility proteins in plant mitochondria": Supplementary_Figures_Huynh_et_al.pptx

### Slide 1
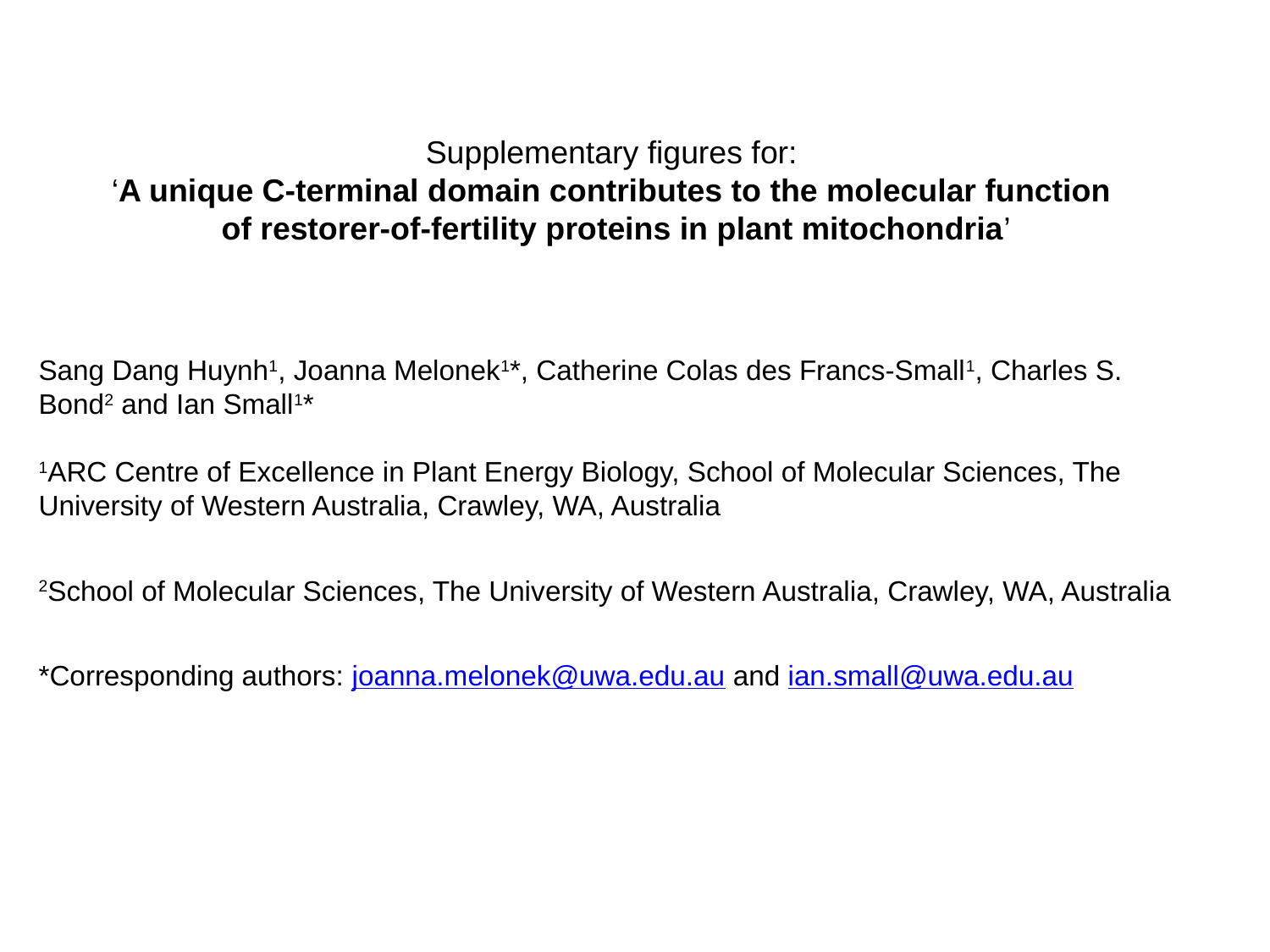

Supplementary figures for:
‘A unique C-terminal domain contributes to the molecular function
of restorer-of-fertility proteins in plant mitochondria’
Sang Dang Huynh^1^, Joanna Melonek^1^*, Catherine Colas des Francs-Small^1^, Charles S. Bond^2^ and Ian Small^1^*
^1^ARC Centre of Excellence in Plant Energy Biology, School of Molecular Sciences, The University of Western Australia, Crawley, WA, Australia
^2^School of Molecular Sciences, The University of Western Australia, Crawley, WA, Australia

### Slide 2
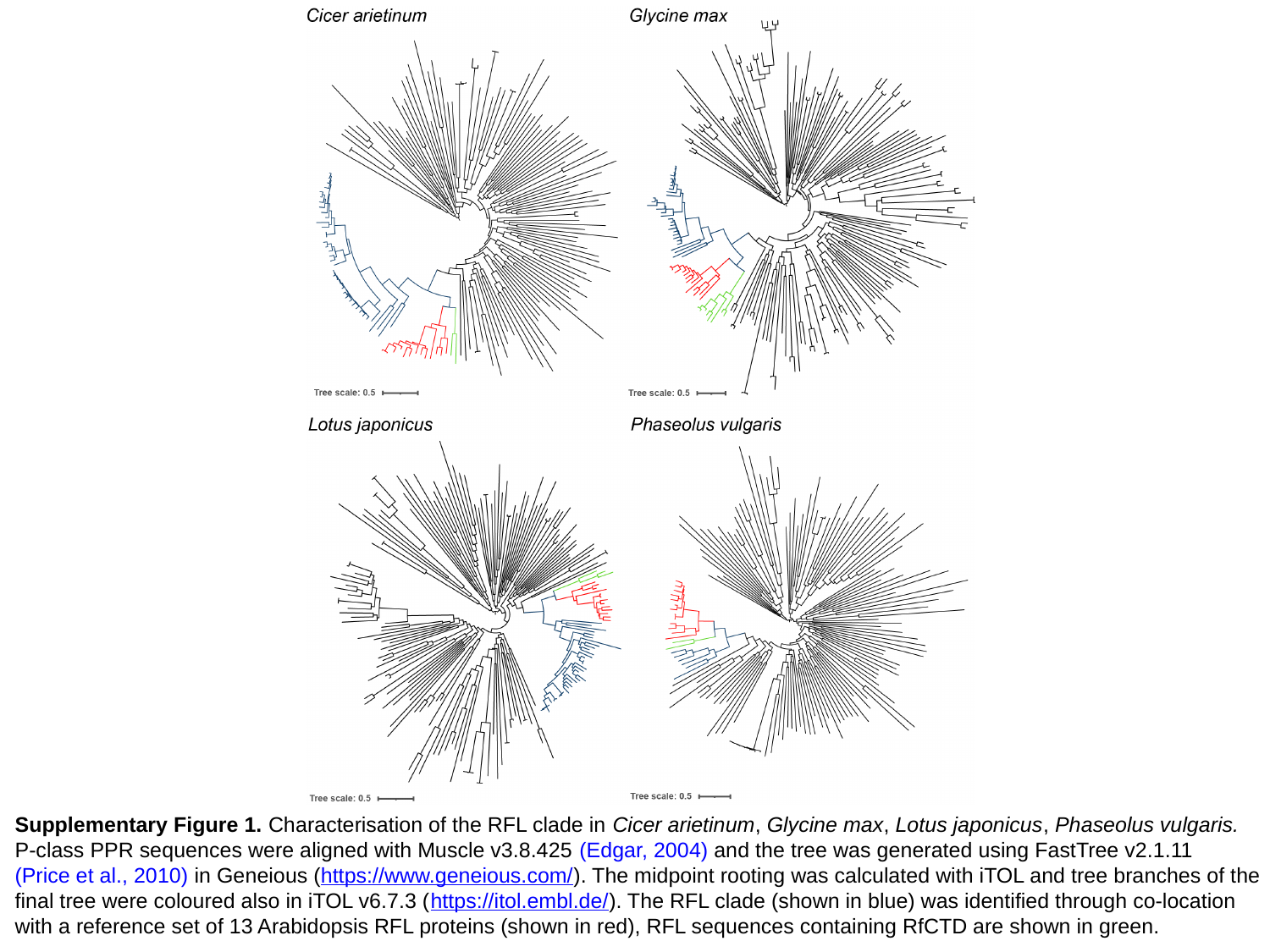

Supplementary Figure 1. Characterisation of the RFL clade in Cicer arietinum, Glycine max, Lotus japonicus, Phaseolus vulgaris.
P-class PPR sequences were aligned with Muscle v3.8.425 (Edgar, 2004) and the tree was generated using FastTree v2.1.11
(Price et al., 2010) in Geneious (https://www.geneious.com/). The midpoint rooting was calculated with iTOL and tree branches of the
final tree were coloured also in iTOL v6.7.3 (https://itol.embl.de/). The RFL clade (shown in blue) was identified through co-location
with a reference set of 13 Arabidopsis RFL proteins (shown in red), RFL sequences containing RfCTD are shown in green.

### Slide 3
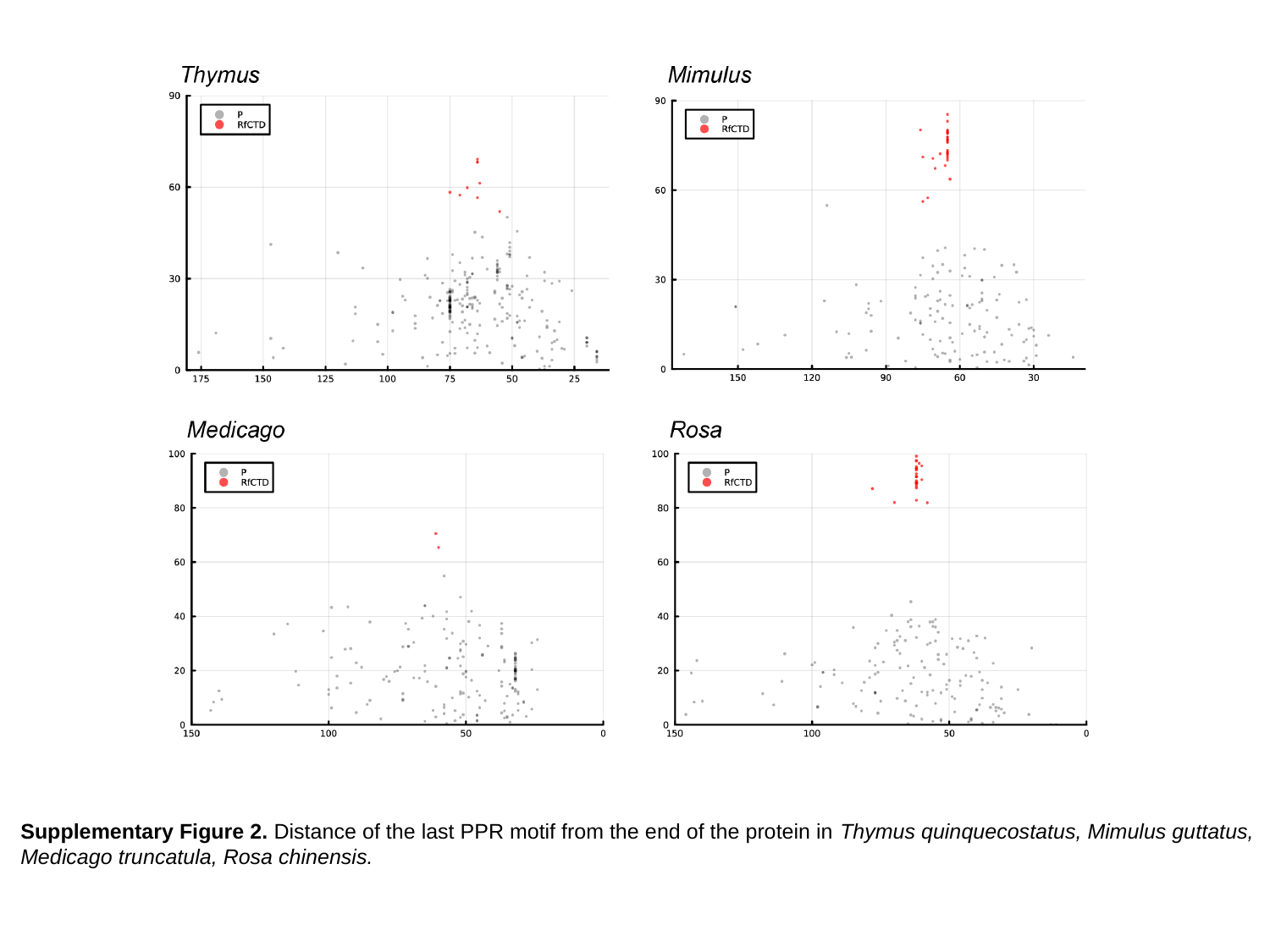

Supplementary Figure 2. Distance of the last PPR motif from the end of the protein in Thymus quinquecostatus, Mimulus guttatus,
Medicago truncatula, Rosa chinensis.

### Slide 4
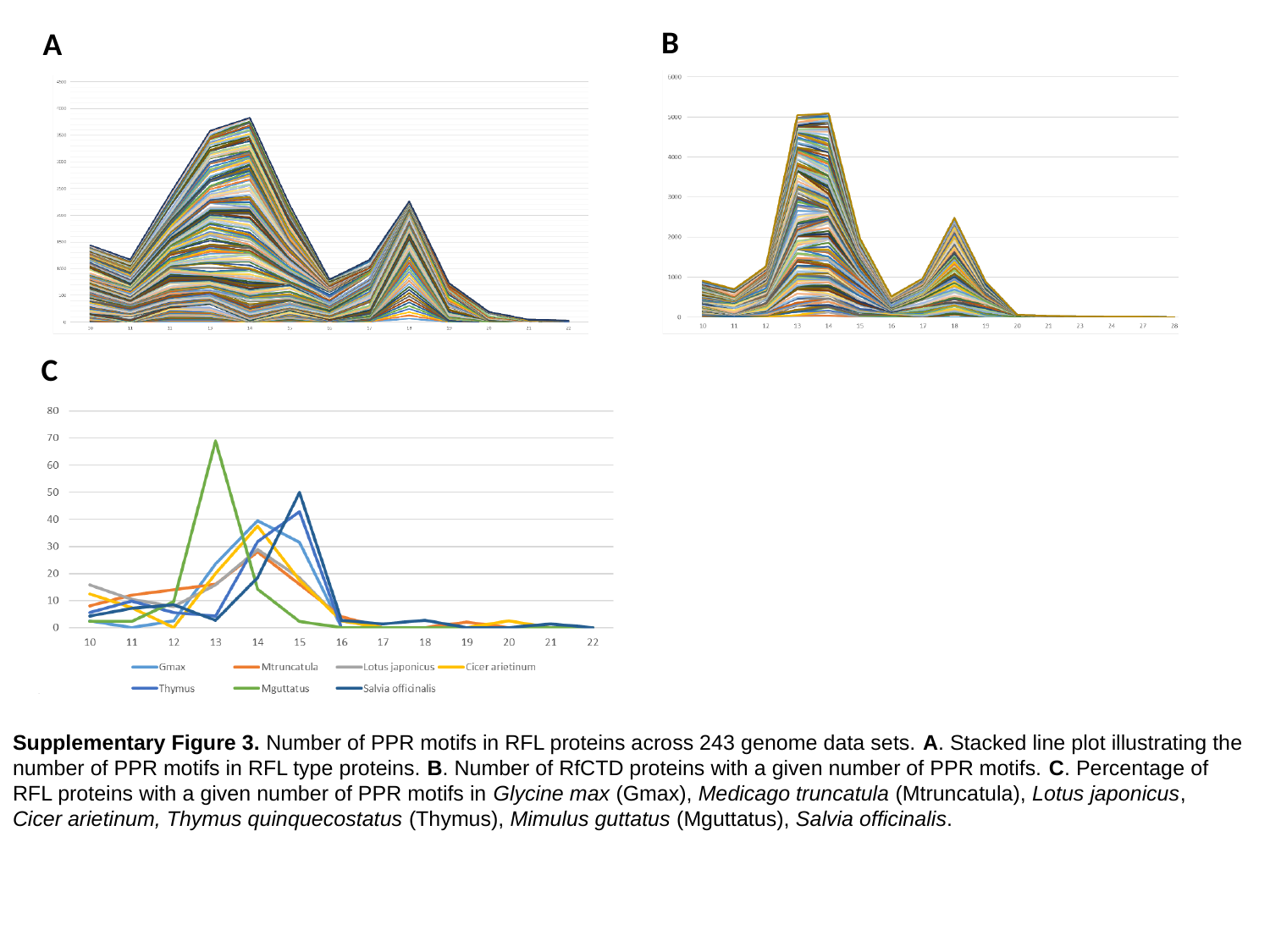

B
A
C
Supplementary Figure 3. Number of PPR motifs in RFL proteins across 243 genome data sets. A. Stacked line plot illustrating the
number of PPR motifs in RFL type proteins. B. Number of RfCTD proteins with a given number of PPR motifs. C. Percentage of
RFL proteins with a given number of PPR motifs in Glycine max (Gmax), Medicago truncatula (Mtruncatula), Lotus japonicus,
Cicer arietinum, Thymus quinquecostatus (Thymus), Mimulus guttatus (Mguttatus), Salvia officinalis.

### Slide 5
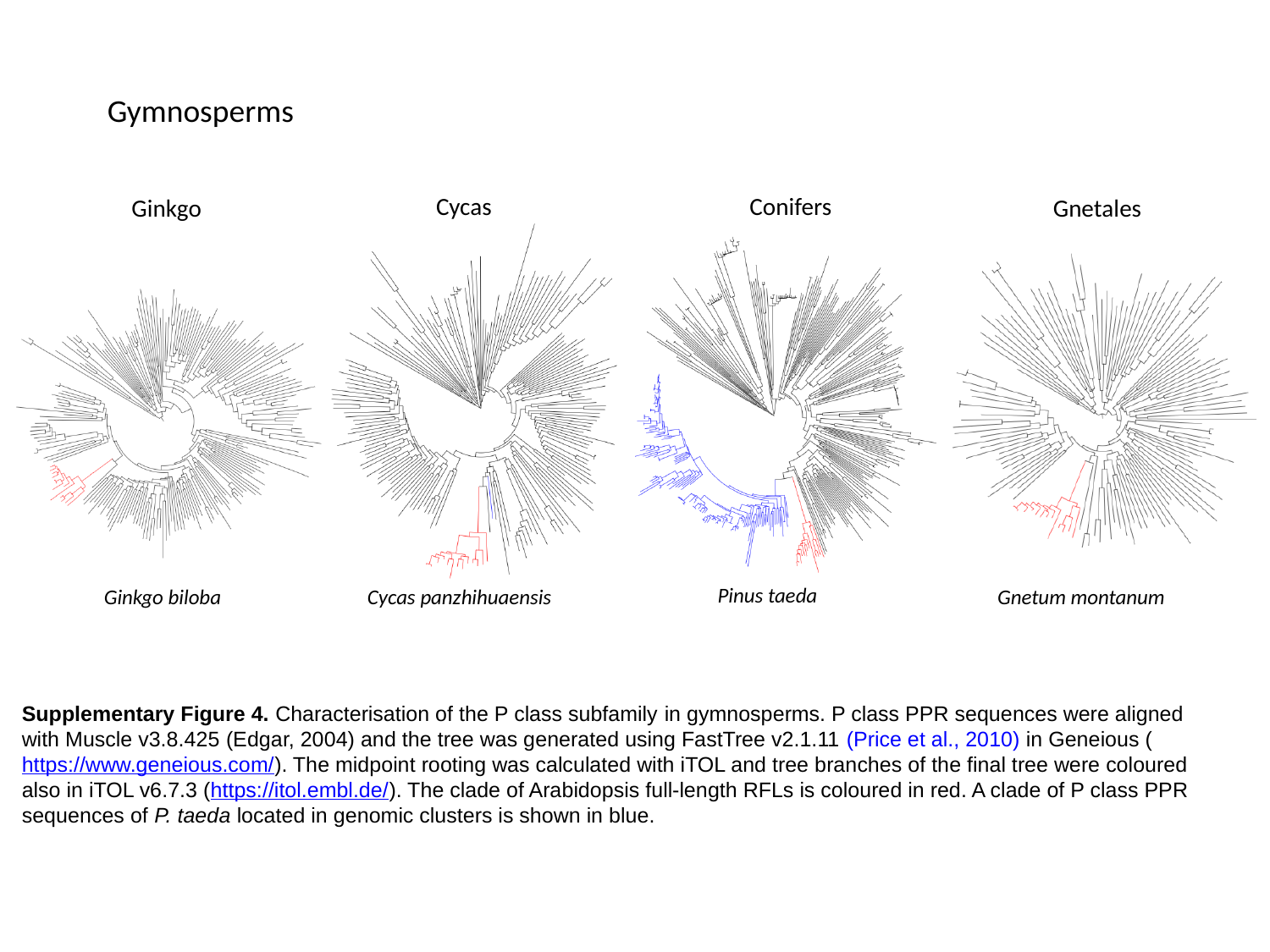

Gymnosperms
Cycas
Conifers
Ginkgo
Gnetales
Pinus taeda
Ginkgo biloba
Gnetum montanum
Cycas panzhihuaensis
Supplementary Figure 4. Characterisation of the P class subfamily in gymnosperms. P class PPR sequences were aligned with Muscle v3.8.425 (Edgar, 2004) and the tree was generated using FastTree v2.1.11 (Price et al., 2010) in Geneious (https://www.geneious.com/). The midpoint rooting was calculated with iTOL and tree branches of the final tree were coloured also in iTOL v6.7.3 (https://itol.embl.de/). The clade of Arabidopsis full-length RFLs is coloured in red. A clade of P class PPR sequences of P. taeda located in genomic clusters is shown in blue.

### Slide 6
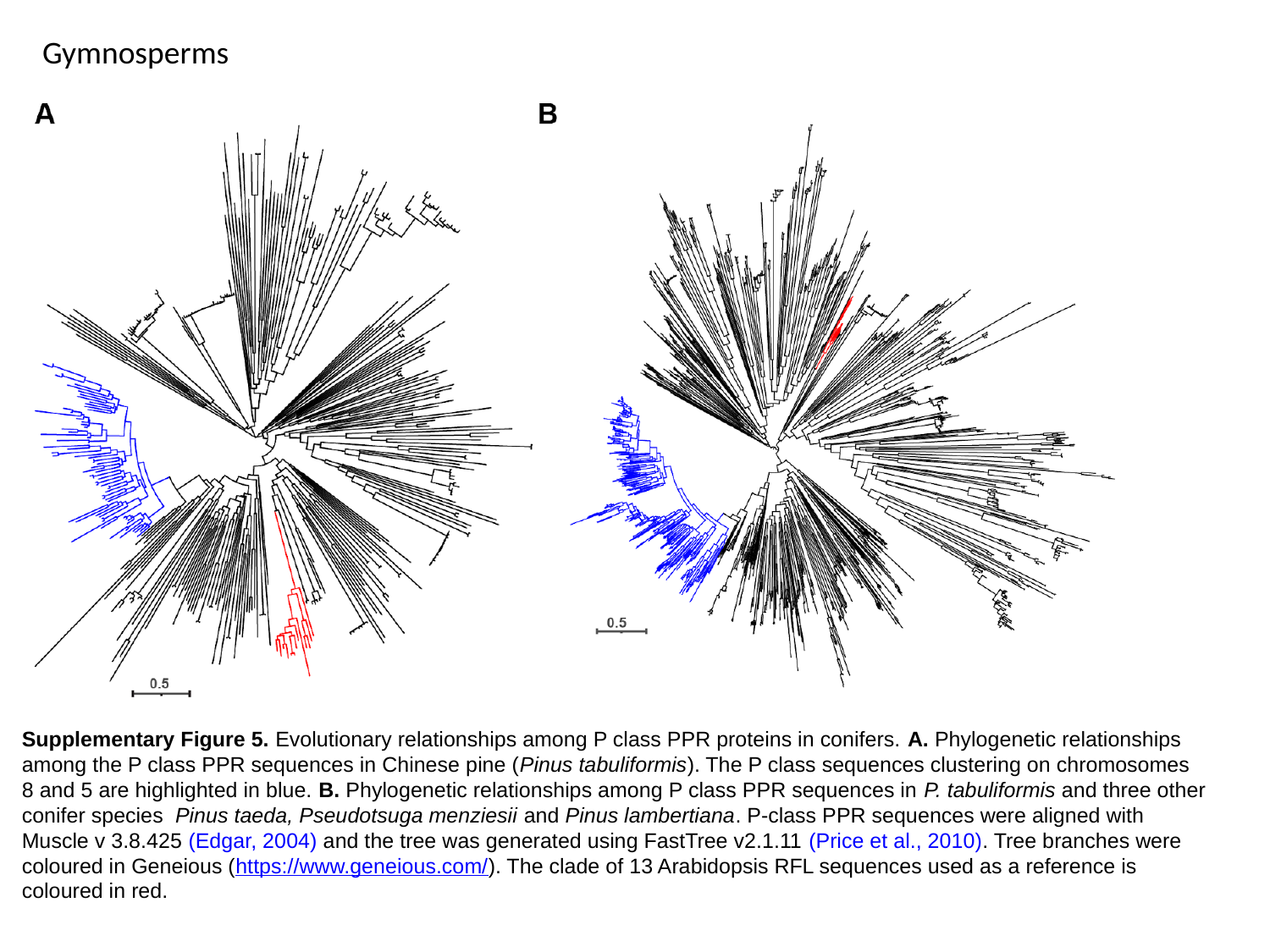

Gymnosperms
Conifers
Supplementary Figure 5. Evolutionary relationships among P class PPR proteins in conifers. A. Phylogenetic relationships among the P class PPR sequences in Chinese pine (Pinus tabuliformis). The P class sequences clustering on chromosomes 8 and 5 are highlighted in blue. B. Phylogenetic relationships among P class PPR sequences in P. tabuliformis and three other conifer species  Pinus taeda, Pseudotsuga menziesii and Pinus lambertiana. P-class PPR sequences were aligned with Muscle v 3.8.425 (Edgar, 2004) and the tree was generated using FastTree v2.1.11 (Price et al., 2010). Tree branches were coloured in Geneious (https://www.geneious.com/). The clade of 13 Arabidopsis RFL sequences used as a reference is coloured in red.

### Slide 7
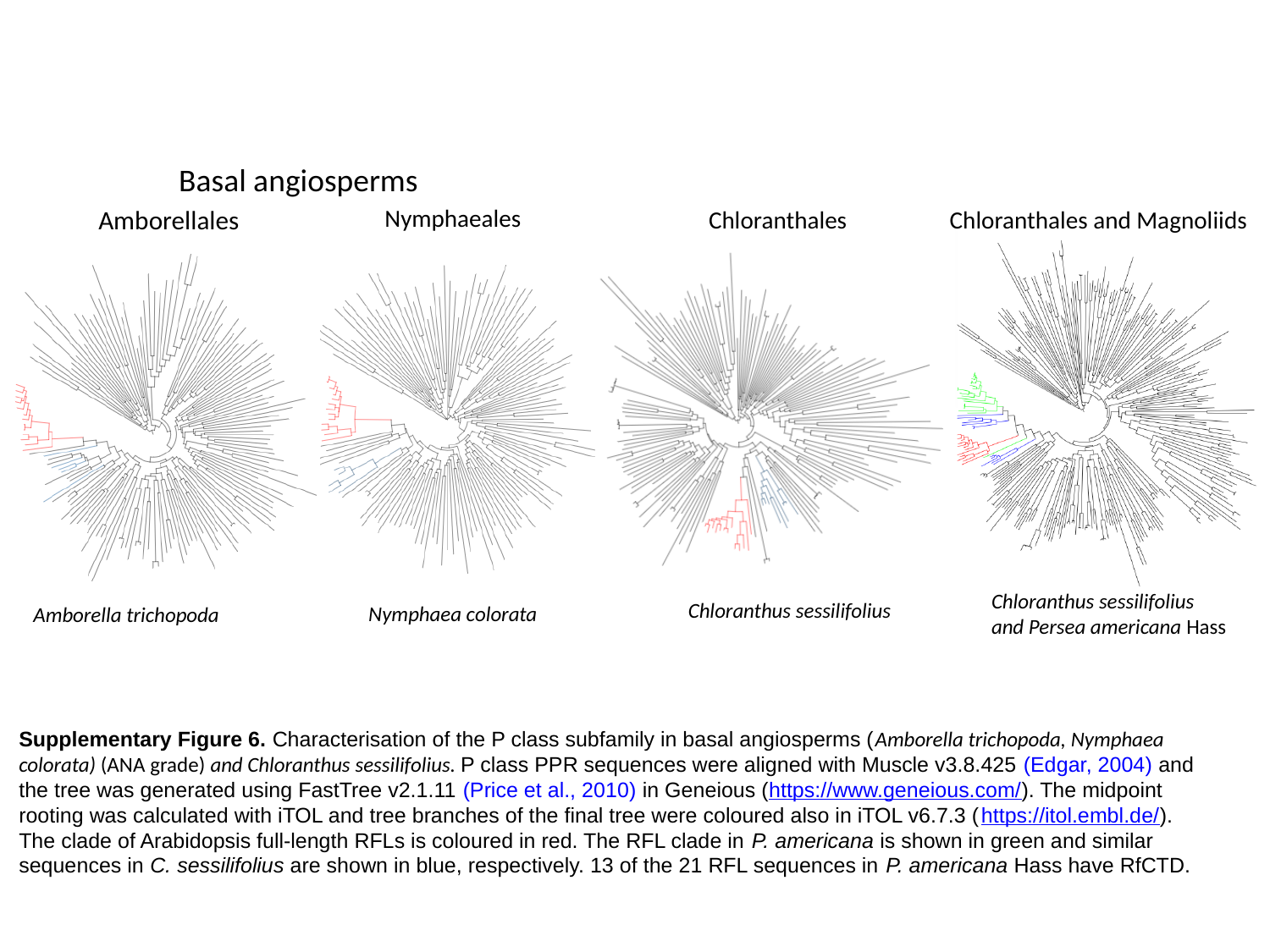

Basal angiosperms
Nymphaeales
Amborellales
Chloranthales
Chloranthales and Magnoliids
Chloranthus sessilifolius
and Persea americana Hass
Chloranthus sessilifolius
Nymphaea colorata
Amborella trichopoda
Supplementary Figure 6. Characterisation of the P class subfamily in basal angiosperms (Amborella trichopoda, Nymphaea colorata) (ANA grade) and Chloranthus sessilifolius. P class PPR sequences were aligned with Muscle v3.8.425 (Edgar, 2004) and the tree was generated using FastTree v2.1.11 (Price et al., 2010) in Geneious (https://www.geneious.com/). The midpoint rooting was calculated with iTOL and tree branches of the final tree were coloured also in iTOL v6.7.3 (https://itol.embl.de/). The clade of Arabidopsis full-length RFLs is coloured in red. The RFL clade in P. americana is shown in green and similar sequences in C. sessilifolius are shown in blue, respectively. 13 of the 21 RFL sequences in P. americana Hass have RfCTD.
